# Harnessing axonal transport to map reward circuitry: Differing brain-wide projections from medial forebrain domains

**DOI:** 10.1101/2023.09.10.557059

**Authors:** E. L. Bearer, C. S. Medina, T. W. Uselman, R. E. Jacobs

**Author notes:** **Correspondance:** Elaine L. Bearer.

## Abstract

Neurons project long axons that contact other distant neurons. Projections can be mapped by hijacking endogenous membrane trafficking machinery by introducing tracers. To witness functional connections in living animals, we developed a tracer detectible by magnetic resonance imaging (MRI), Mn(II). Mn(II) relies on kinesin-1 and amyloid-precursor protein to travel out axons. Within 24h, projection fields of cortical neurons can be mapped brain-wide with this technology. MnCl_2_ was stereotactically injected either into anterior cingulate area (ACA) or into infralimbic/prelimbic (IL/PL) of medial forebrain (n=10-12). Projections were imaged, first by <u>m</u>anganese-<u>e</u>nhanced <u>MRI</u> (MEMRI) live, and then after fixation by microscopy. MR images were collected at 100µm isotropic resolution (∼5 neurons) in 3D at four time points: before and at successive time points after injections. Images were preprocessed by masking non-brain tissue, followed by intensity scaling and spatial alignment. Actual injection locations, measured from post-injection MR images, were found to be 0.06, 0.49 and 0.84mm apart between cohorts, in R-L, A-P, and D-V directions respectively. Mn(II) enhancements arrived in hindbrains by 24h in both cohorts, while co-injected rhodamine dextran was not detectible beyond immediate subcortical projections. Data-driven unbiased voxel-wise statistical maps after ACA injections revealed significant progression of Mn(II) distally into deeper brain regions: globus pallidus, dorsal striatum, amygdala, hypothalamus, substantia nigra, dorsal raphe and locus coeruleus. Accumulation was quantified as a fraction of total volume of each segment containing significantly enhanced voxels (fractional accumulation volumes), and results visualized in column graphs. Unpaired t-tests between groups of brain-wide voxel-wise intensity profiling by either region of interest (ROI) measurements or statistical parametric mapping highlighted distinct differences in distal accumulation between injection sites, with ACA projecting to periaqueductal gray and IL/PL to basolateral amygdala (p<0.001 FDR). Mn(II) distal accumulations differed dramatically between injection groups in subdomains of the hypothalamus, with ACA targeting dorsal medial, periventricular region and mammillary body nuclei, while IL/PL went to anterior hypothalamic areas and lateral hypothalamic nuclei. Given that these hypothalamic subsegments communicate activity in the central nervous system to the body, these observations describing distinct forebrain projection fields will undoubtedly lead to newer insights in mind-body relationships.

## INTRODUCTION

Medial prefrontal pyramidal neurons project long axons deep into distant brain regions. These projections have been mapped by experimentally hijacking endogenous membrane trafficking machinery, axonal transport, that feeds the neuronal synapse by introducing labelled tracers into that transport system, either distally for retrograde transport in the rat (Gabbott et al., 2005), or within the forebrain for anterograde transport in mouse (Fillinger et al., 2018). A variety of clever molecular methods to introduce labelled tracers into specific subgroups of neurons has been developed to map projections from the cortex (Zingg et al., 2014) and other sites. The mouse brain connectivity project at the Allen Institute for Brain Science has mapped projections by fluorescence microscopy in individual mouse brains after fixation including those from medial prefrontal cortex (mPFC) (https://connectivity.brain-map.org/projection) which may add detail to tractography of projections by MRI (Aydogan et al., 2018). Optogenics reveals functional interactions along these traced pathways from forebrain to nucleus accumbens and basolateral amygda (Riga et al., 2014). While there is a long history of results from tract tracing by microscopy, this approach is somewhat limited by the opacity of the brain, which necessitates sacrifice of the animal for subsequent optical imaging, either after sectioning (Fillinger et al., 2018, Gabbott et al., 2005, Sesack and Pickel, 1992) or by rendering the brain transparent for whole brain light-sheet microscopy (Jing et al., 2018). Histologic tracers are typically large molecules labelled either enzymatically, or with an antigen, or with a fluorescent tag. Whether injected or expressed from an exogenous vector, these tracers typically take a long time (weeks) to accumulate distally in sufficient amounts for detection by microscopy.

Tracing projections is critical to our understanding of brain circuitry. To witness functional connections in living animals, we developed a tracer detectible by magnetic resonance imaging (MRI), Mn(II) (Bearer et al., 2007a, Bearer et al., 2007b, Uselman et al., 2022). Initially proposed as an MR contrast agent for axonal tracing by Koretsky’s group (Pautler and Koretsky, 2002), Mn(II) gives a bright signal in T_1_-weighted MRI, enters cells through the voltage-gated calcium channels (Bedenk et al., 2018, Narita et al., 1990), is trafficked in axons by kinesin-based microtubule anterograde transport (Bearer et al., 2007a, Medina et al., 2017a, Pautler, 2006), crosses active synapses (Bearer et al., 2007a, Pautler, 2006, Pautler et al., 2003) and is performed in living animals. Projection mapping by manganese-enhanced MRI (MEMRI) informs on physical connections between neurons, while tractography by diffusion-weighted magnetic resonance imaging informs on the anatomy of axon bundles. Thus, a map of axonal projections emanating from a localized intracerebral injection of MnCl_2_ is acquired by MEMRI. MEMRI has traced projections from nares to olfactory bulb (Pautler and Koretsky, 2002), from retina to visual cortex (Watanabe et al., 2002), from the amygdala to hippocampus (Pautler et al., 2003), from CA3 of the hippocampus to septal nuclei in the basal forebrain (Bearer et al., 2018, Bearer et al., 2007b, Gallagher et al., 2012, Medina et al., 2017a, Medina et al., 2019) and from prefrontal cortex globally into many regions throughout the deeper brain (Bearer et al., 2009a, Gallagher et al., 2013, Zhang et al., 2010) in a wide range of experimental systems. Manganese tract-tracing is particularly useful for diffuse projections, where distal accumulations of Mn(II) can be detected, mapped and intensity values measured as a proxy for connections and relative rates of transport to distal destinations.

Phineas Gage’s personality change after his misadventure in the 1840’s when an iron rod plunged through his forebrain brought attention to this region of the brain in decision making, executive functions and emotions (Damasio et al., 1994). Since then, much progress has been made discovering functional correlates of that region, its projections into the brain, and reciprocal effects of substance use (Frankin TR et al., 2002, Volkow et al., 1996). The nomenclature and segmentation of the medial prefrontal cortex has been under review, with some proposals for a vocabulary that aligns with functional data from other species (van Heukelum et al., 2020). Our nomenclature is based on the Allen Institute Mouse Brain Reference Atlas (Lein et al., 2007) with some correlations to the Paxinos atlas and a more recent update (Vogt and Paxinos, 2014, Paxinos and Franklin, 2001). Our work with MEMRI tract-tracing demonstrated that disruption of three molecular targets of cocaine, SERT, DAT and NET, altered the anatomy of medial prefrontal cortical projections brain-wide (Bearer et al., 2009a, Gallagher et al., 2013, Zhang et al., 2010, Bearer et al., 2009b). Phineas Gage’s injury encompassed a large area in the prefrontal cortex that has been subdivided in more detail by recent experiments. The anterior cingulate and infralimbic/prelimbic areas within the larger medial prefrontal cortex may project to different deeper regions in brainstem, limbic and neuromodulatory structures (Fillinger et al., 2018, Gabbott et al., 2005). These functional connections are not yet fully realized.

Here we take a meta-dimensional approach to map connections from two different regions of the medial forebrain, as detected by Mn(II) accumulations. Our approach differs from previous studies by allowing us to collect and compare data from two sets of animals with each set injected in a different location. After stereotactic intracerebral injections, we collected MRI brain-wide in 4D at 100µm^3^ resolution, the level of a few neurons. Our sample size was large enough to attain statistical significance in unbiased statistical analyses performed on images from each cohort. As the C57BL/6 congenic mouse has minimal inter-individual anatomical differences, this approach produced a map of the average projection anatomy common to all individuals in each cohort. By tracing these forebrain projections brain-wide in a living experimental model, aspects of this controversy may be explored, and differences and similarities between projection anatomies from these two well-defined cortical brain regions revealed. Using an anatomical atlas as pivot to compare between species (Barron et al., 2021), our results may also inform on the human condition.

Using a pre-injection MR image to program a computer-driven stereotactic injection apparatus, we acquired tightly focused intracerebral injection sites, each reproducible across a dozen individuals. Because Mn(II) gives a bright signal, the volume needed for imaging is very small (3-5nL) (Bearer et al., 2022) and produces no detectable permanent damage by histopathology or electrophysiology at this volume and dosage (Uselman et al., 2022). Mn(II) travels quickly, at 5mm/h in the optic nerve (Bearer et al., 2007a), similar to vesicular transport in rat optic nerve (Elluru et al., 1995). Thus MR images captured successively at 6h to 24h report on arrival via transport at distal destinations in living animals (Bearer et al., 2009a, Gallagher et al., 2013, Gallagher et al., 2012, Medina et al., 2017a, Medina et al., 2019, Zhang et al., 2010). By 24h, Mn(II) signal is detected as far as 1.5cm from the injection site, whereas for the histologic tracer, rhodamine dextran, 3 weeks is necessary for detection at much shorter distances and requires sacrifice and fixation to visualize (Bearer et al., 2009a, Gallagher et al., 2013, Zhang et al., 2010). Because the brain is transparent to MRI, the dynamics of transport in the living animal can be followed. As we have shown, Mn(II) transport is in part dependent on the microtubule-based molecular motor, kinesin-1 (Bearer et al., 2007a, Medina et al., 2017a), and on the transmembrane protein, amyloid precursor protein, an adaptor for kinesin-1 (Gallagher et al., 2012, Bearer and Wu, 2019, Satpute-Krishnan et al., 2006). Thus, MEMRI reports on global connections in living mice at relatively short time intervals. Because Mn(II) preferentially crosses active synapses (Bearer et al., 2007a), MEMRI track tracing reveals active trans-synaptic projections.

Computational processing of whole brain image stacks allows sophisticated statistical analyses of large datasets containing hundreds of images from many living individuals, each captured successively at multiple time points. First the brain image is extracted from the living animal’s whole head image (Delora et al., 2016), then all images are normalized, both spatially and intensity-wise, and finally anatomically aligned into a uniform 3D matrix (Medina et al., 2017b). Aligned datasets are subjected to unbiased, voxel-wise statistical parametric mapping and region of interest intensity measurements to map and measure projection patterns and connectivity (Bearer et al., 2009a, Gallagher et al., 2013, Uselman et al., 2020, Zhang et al., 2010). Here we compare distal projections from injections into two different adjacent regions in the medial prefrontal cortex. First, we explore our new dataset obtained after injections into the anterior cingulate area (ACA), and then we compare these results with our archival dataset obtained after injections into the infralimbic/prelimbic region.

## MATERIALS AND METHODS

### Animals

Mice (C57BL/6J) were obtained from JAX, group housed at California Institute of Technology in a climate-controlled mouse house with a 12hr light-dark cycle. Mice were initially imaged after systemic MnCl_2_ by MEMRI according to our protocol (Uselman et al., 2020). Three weeks after the final scan, forebrain projection mapping from the anterior cingulate area was performed. This will ultimately allow us to compare brain-wide neural activity with forebrain projection anatomy. Here we focus on medial prefrontal forebrain projections. All protocols were approved by both Caltech and UNM IACUC. Numbers of mice were determined by a power analysis based on region of interest measurements before and after MnCl_2_. We used all male mice at an average age of 15.5 weeks (14-19 weeks). We previously compared female and male mice and found no statistically significant differences (Uselman et al., 2020). A second dataset of images from mice injected in medial prefrontal cortex that we had previously reported were retrieved from our archives (Bearer et al., 2009b), aligned with this new dataset and processed in parallel. These were all female, ages 19-23 weeks. We followed the same procedures for the new data with ACA injections as we had previously reported for the PL/IL injections except for the injection site locations in ACA vs IL/PL, and the post-injection image collection, where intervals between the first 30m post-injection image and the final 24h image differed(Bearer et al., 2009b). However, to control for the impact of additional prior experience on the new cohort, we compared behavior at baseline, presumably the status of the non-experienced mice in the archival data, and at 3 weeks after experience, when the intracerebral injections for the new data were performed (**Supplementary Fig. S1**). Exploration was video recorded during the last 10m of two 30m periods spent in an arena before (at baseline) and after (23 days, PreFB) experiences just prior to the intracerebral injections. The percent time spent not moving at one-minute intervals was tabulated in Ethovision and spreadsheets of values statistically analyzed in R by ANOVA. We found that at the time point of the intracerebral injection (23 days after baseline and 14 days after the last experiences), there was a minor but statistically significant effect on time spent not moving (p < 0.01) of those manipulations as compared to their baseline.

### MnCl_2_ injection

Both datasets were acquired with the same imaging protocol. Mice were anesthetized in 1.5% isoflurane in room air. After first acquiring a pre-injection image (detailed below), mice were maintained under anesthesia and mounted into the stereotactic frame. After exposing the skull, a small borehole was drilled, with placement controlled by the stereotactic apparatus to avoid injuring the brain, which in mouse is very close to the skull. The drill was then replaced with a micropipette loaded with 3-5nL of 0.6 M MnCl_2_ with 0.3mg/ml 3K rhodamine dextran amine (RDti and 0.15mg/ml 10K fluorescein dextran amine (Molecular Probes/Invitrogen/ThermoFisher) in sterile phosphate-buffered saline. The amount of dextran amine molecules was 0.08 picomoles or approximately 6.9 x 10^16^ molecules. The solution injected into the anterior cingulate area (ACA), with the stereotactic device settings of: x, 0.5 mm right of the midline; y, 1.0 mm anterior to bregma; z, −1.5 deep to the surface of the skull) as previously described (Bearer et al., 2022). Injection proceeded over 5 minutes using a hand-pulled quartz micropipette loaded under direct visualization for precise volume control, guided by computer-assisted stereotactic injector (myNeuroLab.com, IL, USA). After injection, the micropipette was removed slowly, anesthesia was maintained while placement of the mouse into the scanner’s mouse holder with its head secured in a Teflon stereotactic unit to reduce movement artefact. Then the holder was placed into the MR scanner. Within the scanner, anesthesia was maintained (1 – 1.5% isoflurane in medical grade air). Inspection of the first image confirmed injection site placement. Mice received 0.3 ml of saline subcutaneously before each imaging session and again upon removal from the scanner. To prevent drying, eyes were treated with ophthalmic ointment before each scan.

### Image Acquisition

An 11.7 T 89 mm vertical bore Bruker BioSpin Avance DRX500 vertical bore scanner (Bruker BioSpin Inc., Billerica, MA) equipped with a Micro2.5 gradient system was used to acquire all mouse brain images with a 35 mm linear birdcage radio frequency (RF) coil. Temperature and respiration were continuously monitored during data acquisition with temperature maintained at 37°C and respiration maintained at a rate of 100–120 per minute. A 3D multi-echo RARE imaging sequence that has combined T_1_- and T_2_-weighting, with RARE factor of 4, 4 averages, TR/TE eff = 250 ms/12 ms; matrix size of 160×128×88; FOV 16 mm x 12.8 mm x 8.8 mm; yielding 100µm isotropic voxels for a 46m scan time.

Each animal underwent 4 imaging sessions (**Fig. 1**). Images were first captured before MnCl_2_ injection, and then at successive time points, beginning immediately after injection (which we will refer to here as the 30m image) and then at 6h and 24h subsequently. Only small inconsequential variations in the exact time after injection were noted. Between sessions, mice were awakened from the anesthesia and returned to their home cage to move freely with unlimited access to food and water.

**Fig. 1.**
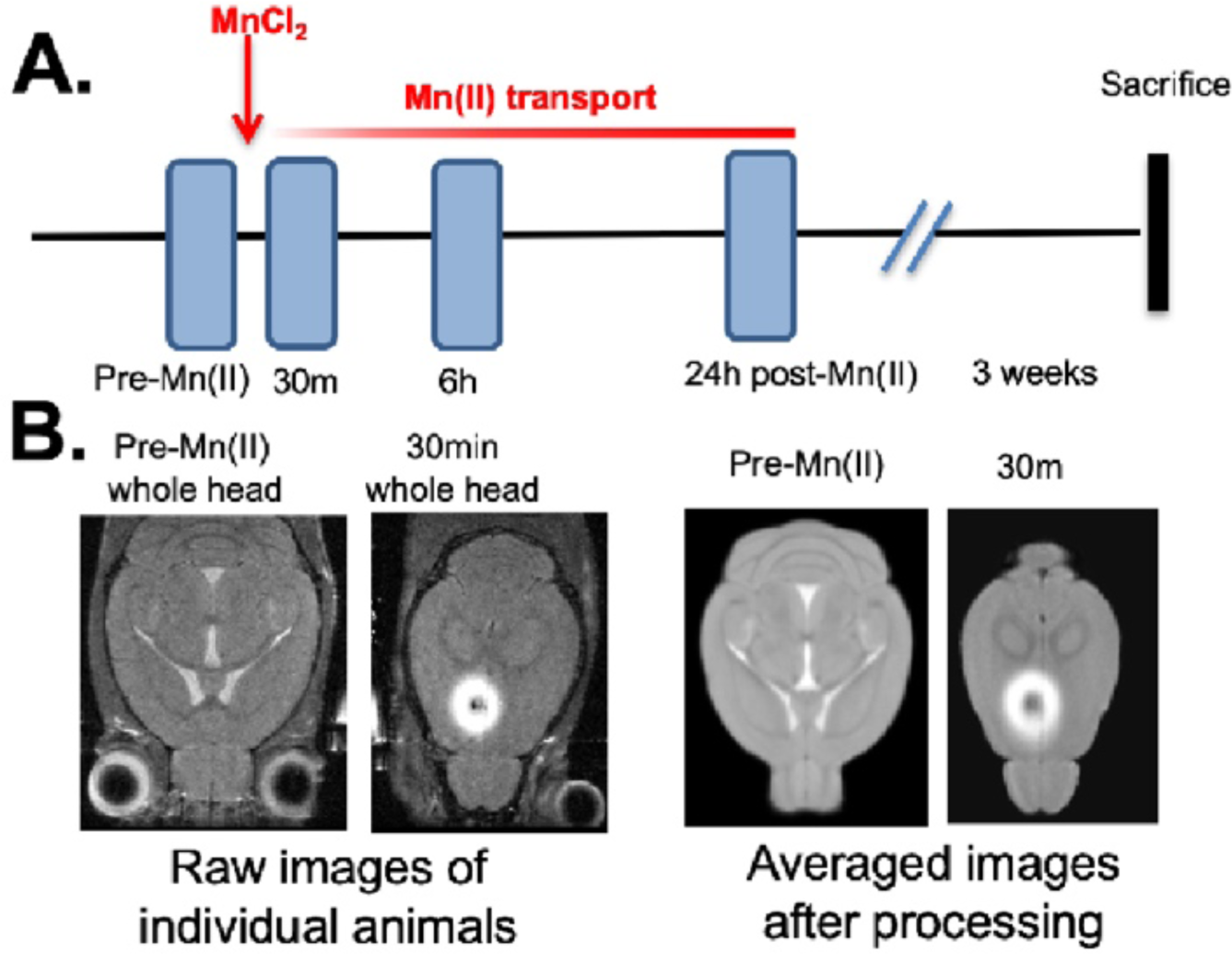
Diagram and examples of experimental procedures. **A)** Timeline of pre-imaging, MnCl_2_ intracerebral stereotactic injection, time-lapse MR scanning and sacrifice for histology. **B)** Slices of MR images at various stages in the processing procedure, from left to right: Raw images of a single animal’s whole head image prior to injection and its 30m post-injection whole head image shown at a higher slice level to pass through the injection site. Both images as taken directly from the scanner and unprocessed. Third and fourth images from right, averaged images from ACA dataset (n = 12) after processing, pre-Mn(II) and 30m post-injection, shown at analogous slice levels as in the first and second raw images to the left respectively. The averaged pre-Mn(II) image represents the MDT. These second set of images have been skull-stripped, N3 corrected, intensity scaled and spatially aligned. All images were captured in 3D at 100 µm isotropic resolution.

### Histology

After the 24h scan, mice were returned to the home cage and at 10-21 days afterwards, euthanized. This delay was to allow for slow transport of the dextran. Each mouse was deeply anaesthetized, and perfused with warm heparinized phosphate buffered saline (PBS; 30 ml) followed by 30ml of room temperature 4% paraformaldehyde (PFA) in PBS. The mouse was decapitated and the head submerged in 4% PFA in PBS overnight at 4°C. Heads were stripped of skin and hair but maintained in the skull at 4°C in 0.01% sodium azide in PBS for three days. After removal from the calvarium, the PBS-azide solution was replenished, and the brains sent to Neurosciences Associates (NSA, Knoxville, TN) for multi-brain embedding, serial sectioning, and staining. Some sections were mounted directly for fluorescence imaging of RDA. Alternate serial sections of the brains were stained for anatomy by thionine (Nissl).

#### Image Normalization

All final images used in this study, both new images reported for the first time here and archived dataset, were processed together in batch. Bruker images were first translated to NIfTI (.nii) files (Neuroimaging Informatics Technology Initiative) and headers matched for all parameters for uniformity of geometry with FSL (FMRIB software library) (Analysis Group, Oxford, United Kingdom) (Lein et al., 2007, Jenkinson et al., 2012a, Smith et al., 2004). Voxel sizes, which were originally captured at 100 µm isopropic, were confirmed using the 3drefit program from AFNI (Analysis of Functional NeuroImages) (Cox, 1996). The brain image was extracted from all non-brain areas within the whole head image by masking (a.k.a. “skull stripping”) according to our standard protocol (Delora et al., 2016). The code for this process is included as Supplemental Data in Delora et al. publication. N3 correction to account for B-field inhomogeneity (Sled et al., 1998, Tustison et al., 2010) was performed on the skull stripped images in MIPAV (Medical Image Processing, Analysis and Visualization) (McAuliffe et al., 2001). Images were intensity normalized using our custom script (Medina et al., 2017b). MATLAB code for this process is included in Supplemental Material with that publication. Our algorithm aligns the modes of each image’s voxel-wise intensity histogram to a single value, calculates and applies the adjustment needed for all other voxels in the image. Here we used one of the IL/PL pre-injection stripped images as the reference for scaling. All images in both datasets were then rigid-body aligned to an image randomly selected from the ACA dataset with the “Realign” function in SPM 12 (Statistical Parametric Mapping) (UCL, London, United Kingdom) (Ashburner et al., 2014, Friston, 1996). Next a minimum deformation transform (MDT) image was prepared from skull-stripped, modally scaled, pre-injection non-contrast-enhanced images from both datasets by a two step-process. First images were aligned to an image randomly selected from within the dataset: here we first selected one from the IL/PL dataset. Resultant images were averaged, creating an averaged image. Then the same set of pre-injection images were re-aligned to that averaged image and resultant images averaged again to create the final MDT. For non-linear alignments, we obtained the control point grid (warp field) from the alignment of the pre-injection image to the MDT. The warp field for alignment of the pre-injection image for each mouse was then applied to that mouse’s post-injection images at all time points after injections using “Normalize” function in SPM 8 (Ashburner et al., 2012). This two-step method avoided potential alignment artefacts that could be introduced during direct warping due to the hyperintense Mn(II) signal at the injection site (Bearer et al., 2009a). While ROI measurements used the final warped aligned dataset, for statistical parametric mapping (SPM) analysis, images were also smoothed with full-width half-maximum (FWHM) Gaussian kernel set to 0.3mm using SPM 12 software package (Statistical Parametric Mapping) (UCL, London, United Kingdom) (Ashburner et al., 2014). Our muse template image (Medina et al., 2017b) was aligned to the MDT using *fsl flirt* (FMRIB software library) (Analysis Group, Oxford, United Kingdom) (Jenkinson et al., 2002, Jenkinson et al., 2012a, Jenkinson et al., 2012b, Smith et al., 2004). For the ACA injection dataset, we also averaged all normalized images at each time point using FSL.

#### Image Analysis

To determine the actual injection site, the hypointense center of the injection site in each animal’s 30m image was identified and its 3D slice position measured in the viewer, FSLeyes, and slice position translated to millimeters relative to midline (ML), bregma (AP), and brain surface. Our planned injection site for dorsal-ventral (DV, depth) included the skull, since we placed the micropipette on the skull surface to program the stereotactic injector. Hence our final location, measured from the surface of the brain and not the skull for the DV dimension is smaller by ∼0.8mm than the planned injection. Average location of injection sites across the dataset was calculated in Excel. Injection sites were then projected onto the 3D minimal deformation atlas rendering of the dataset as “landmarks” in Amira (ThermoFisher).

SPM (SPM 12) (Statistical Parametric Mapping) (UCL, London, United Kingdom) (Ashburner et al., 2014) was used to map Mn(II)-enhancements at 6h and 24h, and to compare maps with archival data from medial prefrontal cortex injections (Bearer et al., 2009b). Original images from our previous study were retrieved and normalized in our pipeline to align with the new images first reported here. These re-aligned images demonstrated that the archival data had injection sites significantly more anterior, in the infralimbic/prelimbic areas (IL/PL), than the current dataset which was in the anterior cingulate area (ACA). We thus first performed within group comparisons on the new data with ACA localized injections. Images at 6h and 24h were each compared to 30m, and 24h to 6h using a paired t-test design in SPM 12. T-values were calculated for p values (0.01-0.0001), uncorrected and with false discovery rate (FDR) corrections for multiple comparisons. We finally chose a p-value threshold based on the most stringent statistic in which some signal was found in all comparisons. We also performed a comparison between 30m and 24h of the archival, IL/PL, dataset by a paired t-test in SPM 12. An unpaired t-test was then performed between 24h images from each experimental cohort. Maps produced by SPM were visualized in FSLeyes and overlaid on gray scale images, either the minimal deformation transform (MDT) of the pre-injected images or our standard “muse_template” aligned to this data. As previously described, our *InVivo Atlas* was hand-drawn on this template based on the Allen Institute for Brain Science Mouse Reference Atlas (Uselman et al., 2020). We determined anatomical locations using our most recently updated *InVivoAtlas* (**Supplementary Table S1** for atlas segments and abbreviations). The atlas consists of three files, a 16bit signed grayscale image at 80 µm isotropic resolution, an 8bit label image at the same resolution and a lookup Table, .lut, which provides the annotations within *fsleyes.* The atlas is first aligned to the MDT using *fsl flirt*, with MDT as input image and grayscale atlas as reference image. A 12-parameter affine transformation of the MDT to the grayscale atlas is performed using Search (incorrectly oriented), Cost function (mutual information), Interpolation (nearest neighbor), and with no weighting. In Mac OS14.2 Terminal, we used the fsl command <convert_xfm> with the inverse .mat file output from flirt to generate an inverse transformation matrix and specify <inverse affine.mat> function. Then we applied the inverse transformation matrix first to the grayscale atlas and then to the annotated labeled atlas in fsl flirt with the <applyxfm>.

### Region of interest (ROI) measurements

We used three interdependent custom shell scripts to drive ROI measurements with fslroi in FSL (Jenkinson et al., 2012a, Jenkinson et al., 2012b, Smith et al., 2004). ROI (3×3×3 voxel cubes) were selected based on the literature as regions receiving input from the mPFC, and coordinates determined based on obvious intensity increases in the average images at 6h or 24h and by statistically significant signal in the SPM. Coordinates for each location were entered into the feeder shell script for automated extraction in FSL. Output values were compiled into a single .csv file which was then loaded into R for statistical analysis using linear mixed model (nlme) and for graphing using ggplot2 (**Supplementary Table S2**, for ROI coordinates).

### Fractional accumulation volumes and column graphs

To quantify the total number of voxels in each segment, our atlas was applied to the template image and the number of voxels within each segment bilaterally calculated in FLS using the *fslmaths* and *fslstats* functions. To determine the number of enhanced voxels at each time point, individual masks for each of the 104 sub-regions identified by the *InVivo Atlas* were generated, also through *fslmaths*. Masks were applied to t-maps produced by SPM, using a statistical threshold of p < 0.05 FDR (within-group) and p < 0.01 FDR (between-group). T-values for this statistic varied: for within group: ACA, 30m > pre (T = 4.056); 6h > 30m (T = 3.912), 24h > 30m (T = 3.573); IL/PL. 30m > pre (T = 4.786) and 24h > 30m (T = 4.809); and for between group: ACA > IL/PL (T = 4.05); and IL/PL > ACA (T = 3.60). The ratios of significantly enhanced to total voxels within each segment were calculated in FSL with the *fslstats* function. These ratios were plotted as column graphs with a customized R script.

### Sub-segmentation of the hypothalamus and identification of bregma slice positions

To define sub-segments within the greater hypothalamus not segmented in our current *InVivo Atlas*, we drew segment boundaries identifiable by grayscale contrasts using the Allen Institute for Brain Science Mouse Brain Reference Atlas, histologic anatomy, as the guide. To determine bregma slice location, we compared anatomy to the Paxinos (Paxinos and Franklin, 2001) using the Online Interactive site (http://labs.gaidi.ca/mouse-brain-atlas/).

## RESULTS

### Uniform injection locations into the ACA, lack of histologic injury at the injection site and entry into expected projections

For this study, we aligned newly acquired ACA injections with archival images from our previous publication on MEMRI tract-tracing from medial prefrontal cortex (mPFC) (Bearer et al., 2009b). We then measured the position of the injection sites in the 30m images of living animals from both cohorts as they appeared in this newly aligned matrix. This allowed quantitative comparisons of injection sites between the two cohorts. The average injection site measured across the immediate post-injection images (30m) from all 12 newly injected animals fell at 0.61 ± 0.11mm right of midline, 0.34 ± 0.27mm anterior to bregma and 0.82 ± 0.18mm deep to the brain surface; whereas average locations within the archival dataset was different (**Table 1 and Supplementary Fig. S2**). This analysis clearly showed that the two datasets had delivered Mn(II) to slightly different locations in mPFC. While both were close to right of midline, the new mPFC injections clustered on average 0.84mm more posterior than those of the archival dataset, which clustered both more anterior and deeper, by 0.49mm. Thus, the archival dataset fell into infralimbic/prelimbic regions (IL/PL), and the new injections, reported for the first time here, clustered into the anterior cingulate area (ACA). We determined that these differences in injection sites could result in delivery of Mn(II) to different groups of mPFC neurons which might project to different distal destinations, as has previously been shown by histologic tract tracing (Fillinger et al., 2018). We thus segregated our data into two cohorts to pursue this idea. Since we had already published some detail from the archival cohort previously (Bearer et al., 2009b), here we first focus on the new data acquired after injections into the ACA.

**Table 1.**
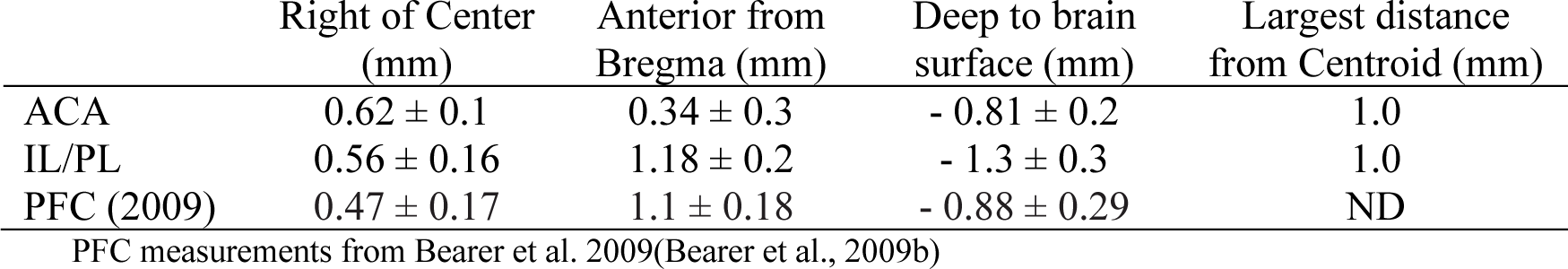
Injection Site Coordinates.

|  | Right of Center<br>(mm) | Anterior from<br>Bregma (mm) | Deep to brain<br>surface (mm) | Largest distance<br>from Centroid (mm) |
| --- | --- | --- | --- | --- |
| ACA | $0.62 \pm 0.1$ | $0.34 \pm 0.3$ | $-0.81 \pm 0.2$ | 1.0 |
| IL/PL | $0.56 \pm 0.16$ | $1.18 \pm 0.2$ | $-1.3 \pm 0.3$ | 1.0 |
| PFC (2009) | $0.47 \pm 0.17$ | $1.1 \pm 0.18$ | $-0.88 \pm 0.29$ | ND |

PFC measurements from Bearer et al. 2009(Bearer et al., 2009b)

Anatomical locations of injections in the ACA dataset were visualized by overlay of our *InVivo Atlas* on an individual animal’s 30m post-injection image (**Fig. 2A**). Mn(II) has a biphasic effect on MR signal, with high concentrations giving a hypo-intense MR signal appearing dark in T_1_-weighted images, and lower concentration giving a bright signal appearing as a halo in this 30m image. Projection of all twelve injection sites as landmarks onto a 3D image of the MDT further demonstrated consistent placement of these injections (**Fig. 2B-C**). Mapping the average location coordinates on the Paxinos online mouse brain atlas (http://labs.gaidi.ca/mouse-brain-atlas/) showed they were within the ACA, labelled as CG1 and CG2 according to Paxinos annotations in this online interactive atlas (**Fig. 2D**).

**Fi. 2.**
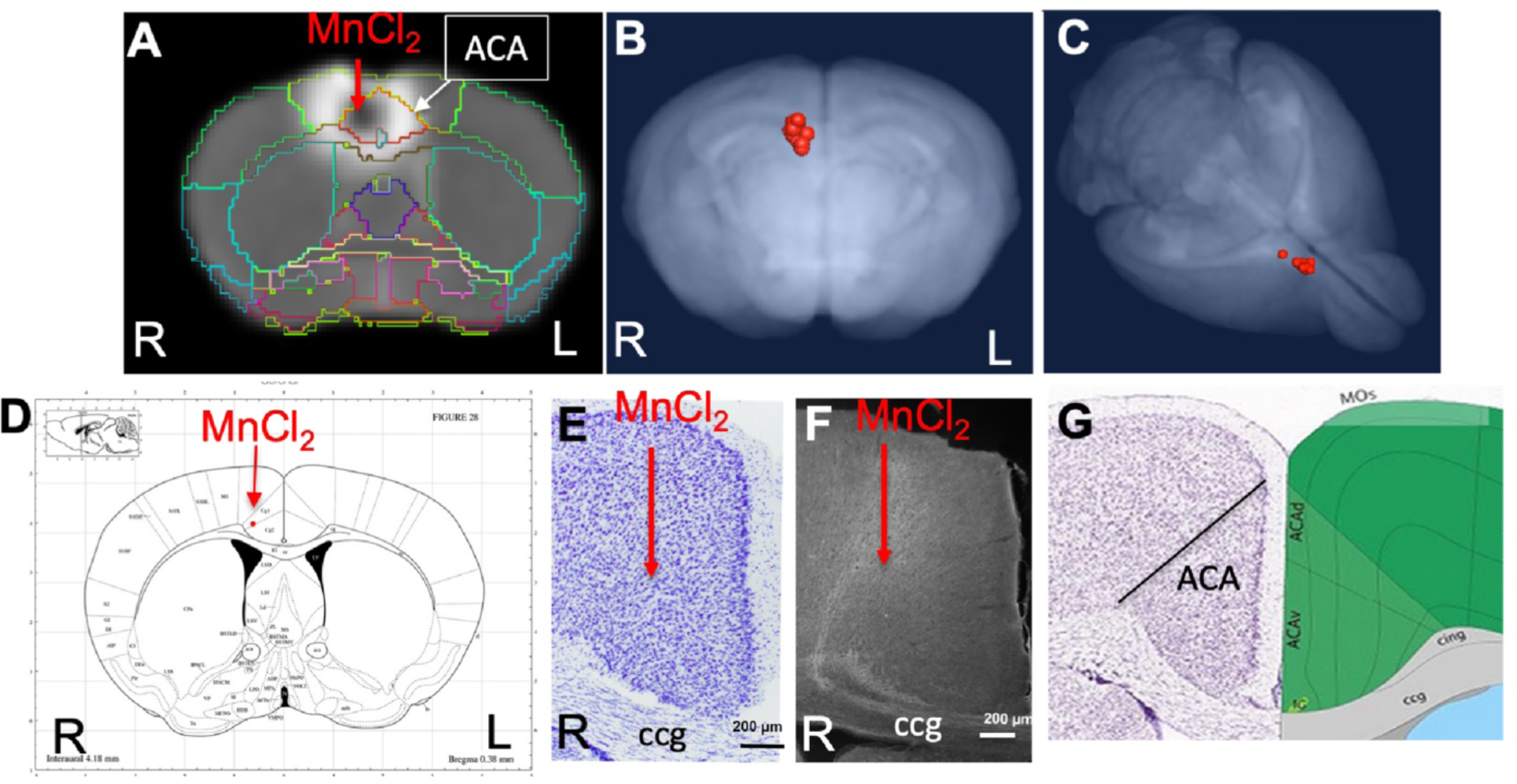
ACA injection site analysis. **A)** Coronal slice from a single animal with our *InVivo Atlas* overlaid. Mn(II) high concentration at center of the injection appears black with a halo of lower Mn(II) concentrations around it. Atlas identifies the injection site in this living mouse as within the ACA. Injection was into the right cortex, with the mouse facing towards the viewer. **B-C)** Center of the injection sites in all 12 mice projected onto a 3D volumetric image of the mouse brain from our muse-template aligned to this dataset. **D)** Screenshot from the online Paxinos atlas (http://labs.gaidi.ca/mouse-brain-atlas/) with average coordinates from the MnCl_2_ injection site analysis placed onto the atlas as a red dot. These coordinates appear precisely as in our digital *InVivo Atlas*, within the ACA which in this atlas is labelled Cg1 and Cg2 for cingulate gyrus. **E)** A section through the ACA from a single mouse in this dataset stained by Nissl. No injection site can be identified. Corpus callosum (ccg). **F)** Rhodamine fluorescence identifies neuronal processes emanating out of the ACA with no clear injection bolus. Projection go both into the cortex and down into fiber tracts of the corpus callosum (ccg) and cingulum bundle (Bubb et al., 2018). **G)** A screenshot of same region segmented and annotated from the Allen Institute Mouse Brain Reference Atlas. The ACA is sub-segmented and both ccg and cingulum bundle (cing) labelled (Lein et al., 2007).

The injection site was not detectable histologically, as no evidence of long-term injury could be found in thianine-nissl-stained serial sections from animals fixed 2-3 weeks after injection, as we have previously reported (**Fig. 2E**) (Bearer et al., 2009b). Only by searching for co-injected dextran fluorescence across serial sections were injection sites identified histologically (**Fig. 2F**). RDA was identified in a region consistent with the ACA, with fluorescent tracer transported along apparent axons into corpus callosum (cc) and cingulum bundle (cing), regions identified according to visual comparisons with the Allen Institute Mouse Brain Reference Atlas (**Fig. 2G**) (Lein et al., 2007). These local processes resembled those reported in individual mice traced by fluorescent tracers in the connectome project by Allen Institute for Brain Science. Microscopic evaluations of the ACA-injected brains also attested to minimal injury and accurate placement into the ACA.

### Mn(II) enhancements visualized in averaged 3D images after injection into the ACA

To ascertain that Mn(II) had successfully progressed into the brain, we averaged the aligned images from each time point and inspected coronal slices for intensity increases in expected locations. Whole brain images captured in living animals by time-lapse MRI were aligned to the pre-injection MDT and then the subset of images for each time point (pre-injection and 30m, 6h and 24h post-injection) across the 12 animals were averaged (**Fig. 3**). Here we show coronal slices from these averaged 3D images at each of the 4 time points, with slice position selected to pass through expected target locations of ACA projections. Mn(II) enhancements were readily apparent in these slices and followed expected ACA projections: dorsal striatum (DS), reticular nucleus of the thalamus (RTN), and substantia nigra reticularis (SNr). In this inspection, we also see the differences in time at which enhancement appears in each of these distal targets, in ipsilateral DS at 6h and in both ipsilateral and contralateral DS at 24h, in the contralateral RTN at 6h where it then fades at 24h, and finally reaching SNr by 24h, possibly bilaterally. Also apparent is the decrease of Mn(II) intensity over time in the injection site, where it virtually disappears by 24h. These visual inspections are not sensitive enough to detect all the projections of the ACA, the dynamics of accumulations between time points, or the degree or statistical significance of signal enhancements.

**Fig. 3.**
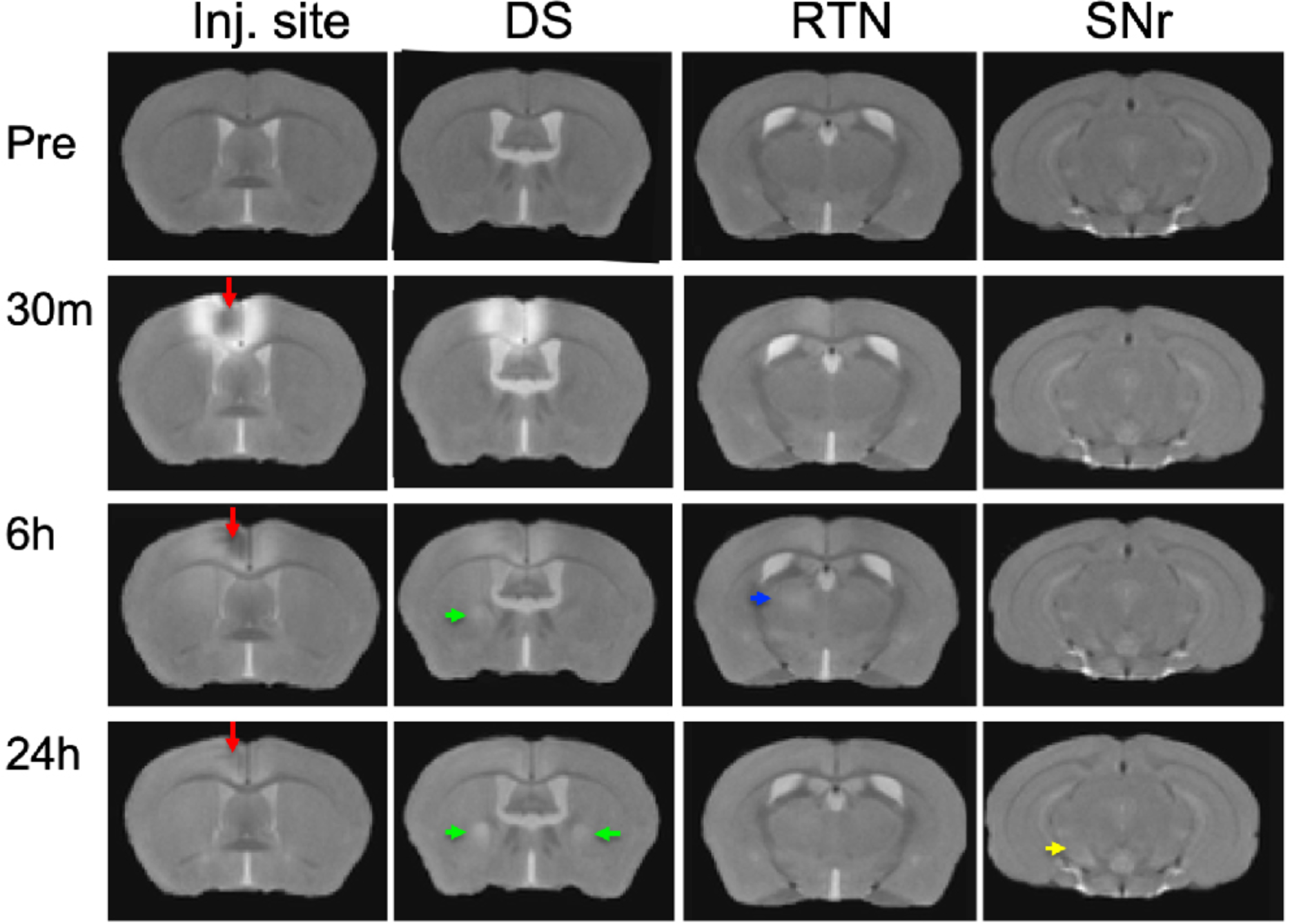
Mn(II) enhancements appear in multiple regions across slices at each time point. Coronal slices were selected at four anterior-posterior locations from averaged aligned 3D images of the twelve animals at each time point. Arrows indicate hyperintense signals: Red, injection site; Green, dorsal striatum (DS); Blue, reticular nucleus of the thalamus (RTN); and Yellow, substantia nigra reticulata (SNr). Anatomy was confirmed by reference to our digital *InVivo Atlas*.

### Statistical parametric mapping of Mn(II) intensity enhancements after ACA injection

To increase sensitivity and identify regions brain-wide that received significant Mn(II) after localized injections in the ACA at each progressive time point, we adopted statistical parametric mapping (SPM) (Bearer et al., 2007b). Such computational approaches detect statistically significant voxel-wise intensity changes in an unbiased and brain-wide comprehensive manner which has broken open our ability to understand human brain dynamics from BOLD MR imaging. When applied to MEMRI, SPM reveals where the Mn(II) has had a significant impact on the MR signal (**Fig. 4**). SPM shows that Mn(II) enhancements progressed from the injection site into the contralateral cortex, as could be seen in MR images, and traveled deeper into subcortical regions. At 6h, shown in red, statistically significant Mn(II) signal occurred in the DS and tracked down, as seen best in the sagittal slice outlined in white, trafficking all the way to the SNr in the brainstem. From 6h to 24h, Mn(II) signal transported even deeper into these structures and also appeared in the retrosplenial cortex and dorsal hippocampus. While this cortical signal could be dismissed as diffusion from the injection site, the lack of signal at 6h (red) between the 30m signal (green) at the injection site and the 24h (blue) suggests that the 24h signal in the retrosplenial cortex is at least partly produced, not by diffusion from the injection site, but by transport either along the corpus callosum from one cortical region to another or indirectly from a polysynaptic circuit first to DS or RNT and then back up to the cortex progressing to the hippocampus.

**Fig. 4.**
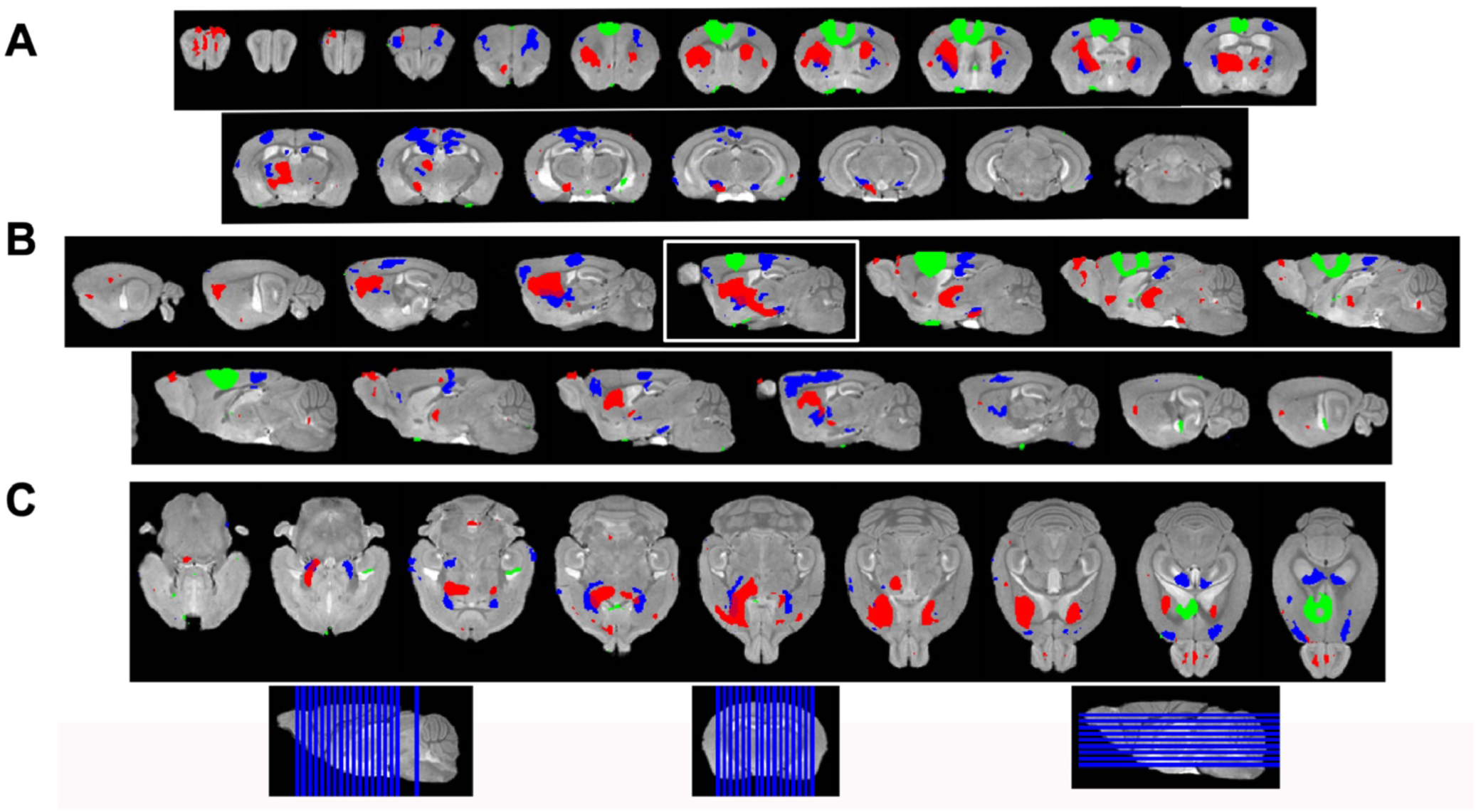
Statistical parametric mapping shows progress of Mn(II)-dependent intensities across time points. **A)** Coronal slices of SPM maps from each time point overlaid on the high definition template shown left to right in the figure from anterior to posterior at 0.5mm intervals. **B)** Sagittal slices at 0.5mm intervals taken from slices right to left side of the brain shown going left to right in the figure. **C)** Axial slices from ventral to dorsal at 0.5mm intervals, shown left to right in the figure. Green, 30min > pre-Mn(II); Red, 6hr > 30m; Blue, 24h > 30m. Slice positions diagrammed at bottom of the figure on the whole brain 3D image.

We then took two additional steps to parse the segmental distribution of transported Mn(II), and thereby obtain clues about ACA projection anatomy: 1) Measuring intensity increases with region of interest analyses; and 2) Quantifying the volume of enhanced voxels, the fractional accumulation volume, in every segment across the brain by applying our *InVivo Atlas* to enumerate statistically enhanced voxels and obtain the ratio of enhanced to total voxels. Finally we moved on to compare fractional accumulation volumes, the volume of Mn(II) enhancement within each segment, between ACA and the IL/PL injection sites with two-tailed unpaired statistical models.

### Mn(II) transports along predicted pathways after ACA injection by region of interest analysis (ROI)

To quantify Mn(II) distal accumulations over time in the ACA injection group, we selected 8 cubic regions of 3×3×3 voxels bilaterally which were predicted by previous reports to receive input from ACA, and measured average intensity values (**Fig. 5**). Measurements were made in 30m, 6h and 24h images across the ACA dataset. Segments were identified with our *InVivo Atlas*. Later time points (6h and 24h) were determined as a ratio of signal intensity in that same ROI divided by intensity in the 30m image. We reasoned that any signal distal to the injection site at 30m after injection was more likely due to rapid interstitial diffusion of the Mn(II) than to slower membrane-bound vesicular transport (see **Supplementary Tables S2** for ROI coordinates, **Table S3** for measurements and **Table S4** for statistics).

**Fig. 5.**
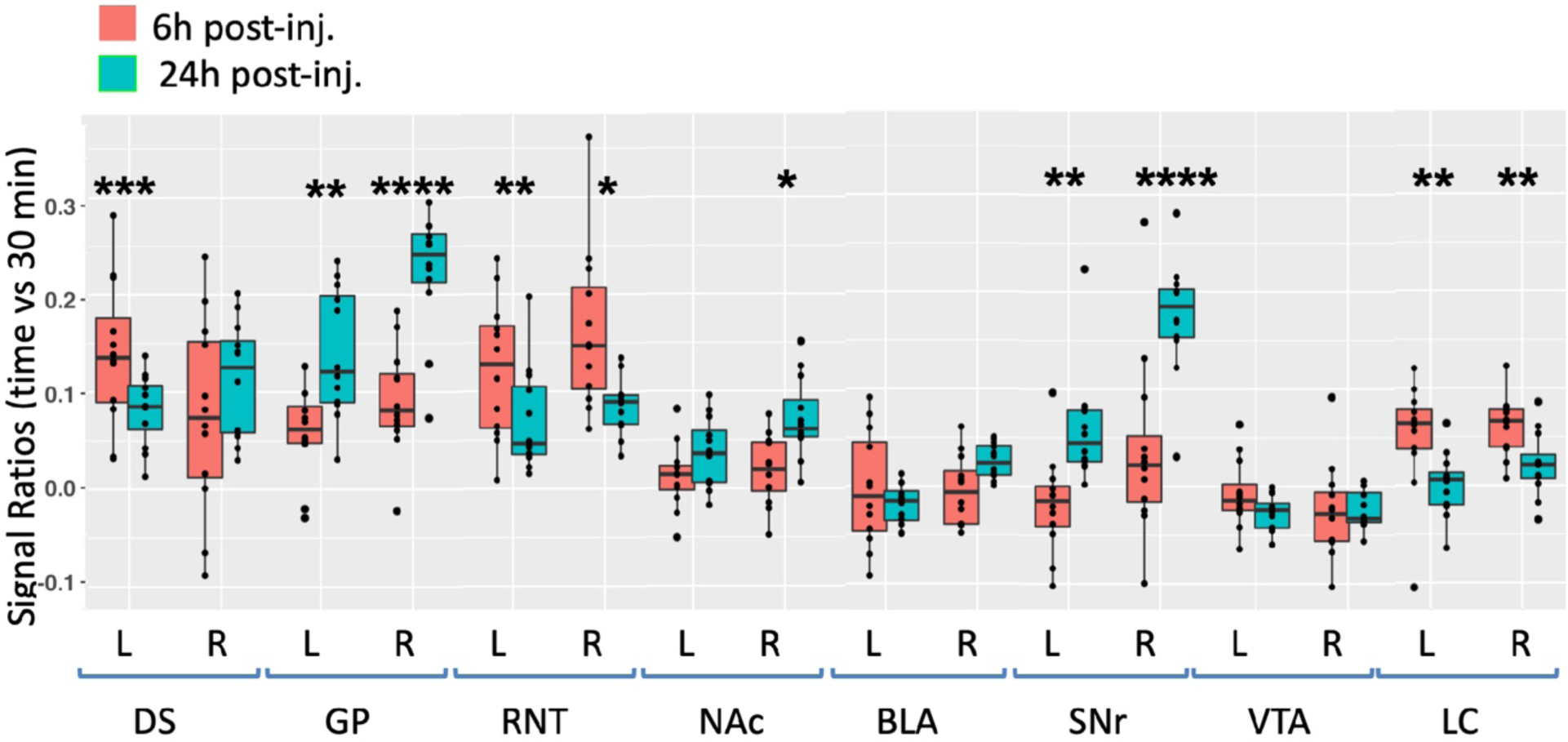
Degrees of intensity changes in regions of interest (ROI) over time after ACA forebrain injection. Measurements of intensity values in 3×3×3 voxels selected in dorsal striatum (DS), globus pallidus (GP), reticular nucleus of the thalamus (RTN), nucleus accumbens (NAc), basolateral amygdala (BLA), substantia nigra reticulata (SNr), ventral tegmental area (VTA) and locus coeruleus (LC). Both contralateral (left, L) and ipsilateral (right, R) were measured in the 30m, 6h and 24h images. Intensity values were calculated as a ratio at 6h (orange) or 24h (green) over that of the 30m images. Box and whisker plots are shown with individual values projected as scatter plots. Some median values increased between 6h and 24h and other decreased. Outliers greater than 2 standard deviations are shown in the scatter plot and included in the statistics. Statistical significance of comparisons by linear mixed model between 6h and 24h intensities regardless of directionality as indicated: * p < 0.01; ** p <0.01; *** p < 0.0001, **** p < 0.00001.

At 6h after ACA injection, signal increased more than 10% in both left and right DS and RNT, and on the right GP, while the LC bilaterally had > 5% intensity increases. We compared increases from 6h to 24h statistically and found significant increases in both sides of GP and SNr, and in the right NAc ipsilateral to the injection site. A small increase of a few percent was also detected in the right BLA at 24h, which was not statistically significance. These increases likely represent continued transport to these regions, possibly secondarily from accumulations at other targets within a polysynaptic projection anatomy. In contrast to the increasing signal at 24h in these ROIs, signal decreased significantly in a few others compared to the 6h time point, with decrease on both sides of RNT and LC. These decreases compared to the loss of signal in the injection site shown in **Fig. 3** may represent less Mn(II) entering the transport system from projections that directly synapse on these segments at later time points.

### Projection fields differ in ACA injection compared to IL/PL

To pursue the possibility that these minor differences in forebrain injection sites might trace different anatomical projections, we first performed voxel-wise paired t-test by SPM on each aligned dataset separately and compared results between groups by visual inspection (**Fig. 6, upper panels**). Differences in the placement of injection sites were obvious in sagittal slices with 30m time point images overlaid: ACA shown on the left panel in green, and IL/PL on the right in yellow (also see **Supplemental Fig. S2**). These contrasts also appear in the fractional accumulation volumes shown in the two within group column graphs (**Fig. 6, lower panels**). In segments within the injection site in the ACA cohort, there was a large ratio (0.7) of enhanced to total voxels in the ACA segment, and only a low ratio of enhanced to total voxels within IL/IP segments (**Fig. 6, lower panels, arrows**). In contrast, in the IL/PL cohort, the ACA segment has less volume occupied by statistically enhanced voxels (0.4), and both IL/PL segments had 0.6 ratio of enhanced voxels to total. At 6h in the ACA cohort (**Fig. 6, left panels, red**), SPM maps showed progression of significantly enhanced voxels down through the brain stem as far as the SNr with a track that appears fairly diffuse in both groups. Column graphs at 6h and 24h time points in the ACA group show fractional accumulation volumes increased in some but decreased in other segments between 6h and 24h. Since we only have 30m, 1h40m through 4h20m and the final 24h post-injection images for the IL/PL group, we could only compare 30m and 24h images between the groups (Bearer et al., 2009b).

**Fig. 6.**
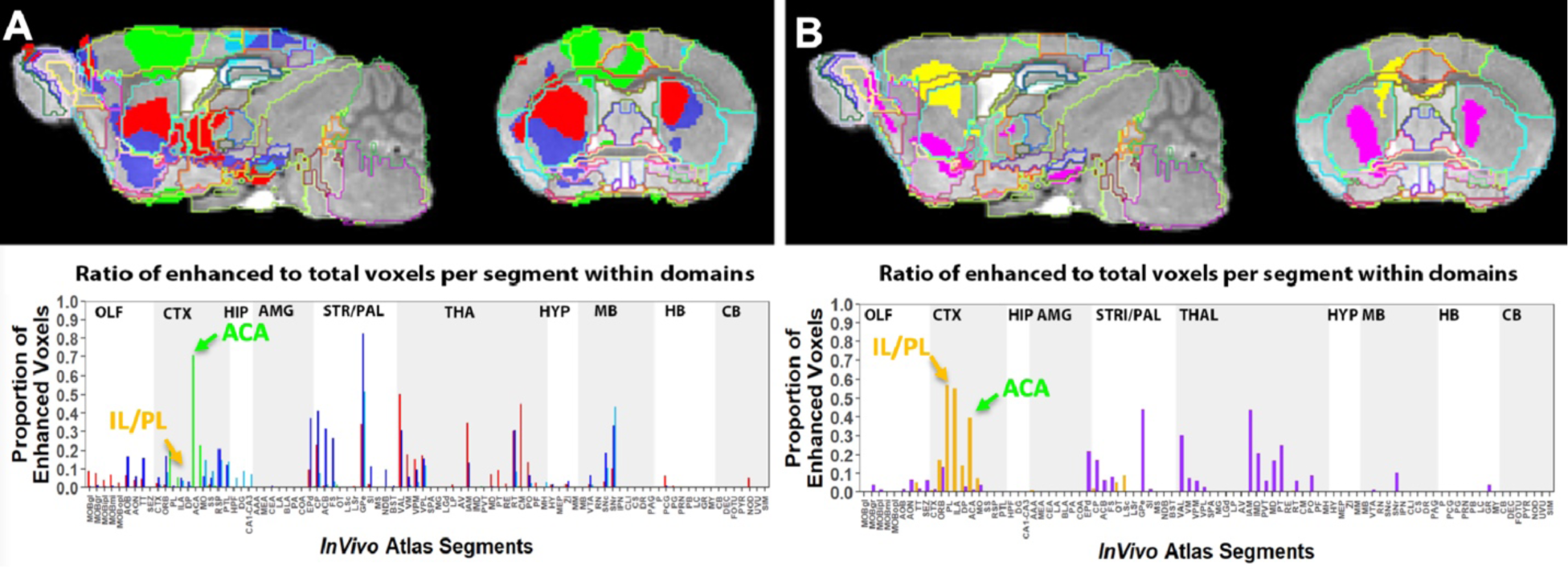
Comparison of projection maps from ACA with IL/PL by visual inspection of within group statistical maps and segment-wise volumetric column graphs. **A)** Slices from voxel-wise brain-wide maps of the anatomy of statistically enhanced voxels after ACA forebrain injection projected onto a grayscale image of our high definition template are shown: Green, 30m > pre-Mn(II) (T = 4.05); Red, 6hr > 30m (T = 3.91); Blue, 24h > 30m (T = 3.57); Turquoise, 24h > 6h (T= 3.97) (p < 0.05 FDR for all comparisons). In this overlay, images are layered on the template in the following order: 24h > 6h, 24h > 30m; 6h >30m; 30m > pre-Mn(II) with the *InVivo Atlas* v10.2 segmentation delineated on the top layer. **B)** Similar overlays but for the IL/PL injections, with Yellow, 30m > pre-Mn(II) (T = 4.78); Pink, 24h > 30m (T = 4.8) (p < 0.05 FDR for both).

Comparisons of statistical maps and fractional accumulations volumes between groups at 24h showed Mn(II) enhancements progressed further in the ACA than in IL/PL group (**Fig. 6, left panels, blue compared to right panels, purple**), with greater accumulation and deeper progression in the ACA cohort. In contrast, in the IL/PL cohort, there was less progress into the midbrain (MB) or hindbrain (HB) segments at 24h. These SPM and column graphs of fractional accumulation volumes extend the information gleaned from ROI analysis by detecting subsegmental dynamics and identifying additional regions not previously recognized and thus not selected *a priori* as candidate regions. Such additional segments in the ACA cohort include APB and AON in the olfactory system, EPd in the amygdala, CP and FS in the striatum/palladum, RSP in the cortex, VAL, IAM and CM in the thalamus, and small volumes of statistically enhanced voxels in the hind brain (PCG) and cerebellum (NOD).

In comparison, the IL/PL injected cohort seemed to have less distal accumulation, suggesting lower transport overall, although the total number of enhanced voxels in the forebrain was similar in both cohorts testifying to equivalent amounts of Mn(II) was injected. Differing distal destinations could obscure some accumulations in the IL/PL as segment volumes are not uniform. Although small alterations in these experimental procedures between individuals within the cohorts could have inadvertently occurred, any effect from this possibility would be minimized by our sample size, 10-12 animals in each cohort, and statistical analyses. While IL/PL were all female, the ACA were all male, partly due to changes in IACUC regulations, although both groups were the same strain and similar ages. The ACA mice had also undergone additional manipulations while at Caltech not experienced by the IL/PL injected cohort.

Many similarities and a few differences in segmental accumulations appeared in visual comparisons between segment-wise column graphs. A most obvious difference in addition to the injection site was more transport to the midbrain after ACA injection and possibly more to the thalamus after IL/PL injection. These visual inspections were not as satisfying as a direct statistical comparison by unpaired t-tests between the two groups.

### Between group analyses reveal dramatic, statistically significant differences in distal accumulations from different forebrain injection locations

For a direct comparison between groups, we first measured the same ROIs as described in **Fig. 5** in both ACA and IL/Pl groups (**Fig. 7**). We picked the 24h time point to compare, as that was captured at nearly identical times after injection in both groups. We calculated the ratio of average MR intensity signal at 24h versus 30m in each group and performed statistical comparisons between groups. In four of the eight segments, measured bilaterally, the difference between groups was significant, with ACA greater than IL/PL in DS bilaterally and in ipsilateral (right) SNr, and with IL/PL significantly greater than ACA in contralateral (left) RNT and marginally significantly increased in left BLA as compared to left BLA in the ACA. These results led us to perform a voxel-wise brain-wide unpaired t-test between the two groups with SPM and then calculate fractional accumulation volumetric differences (**Fig. 8**).

**Fig. 7.**
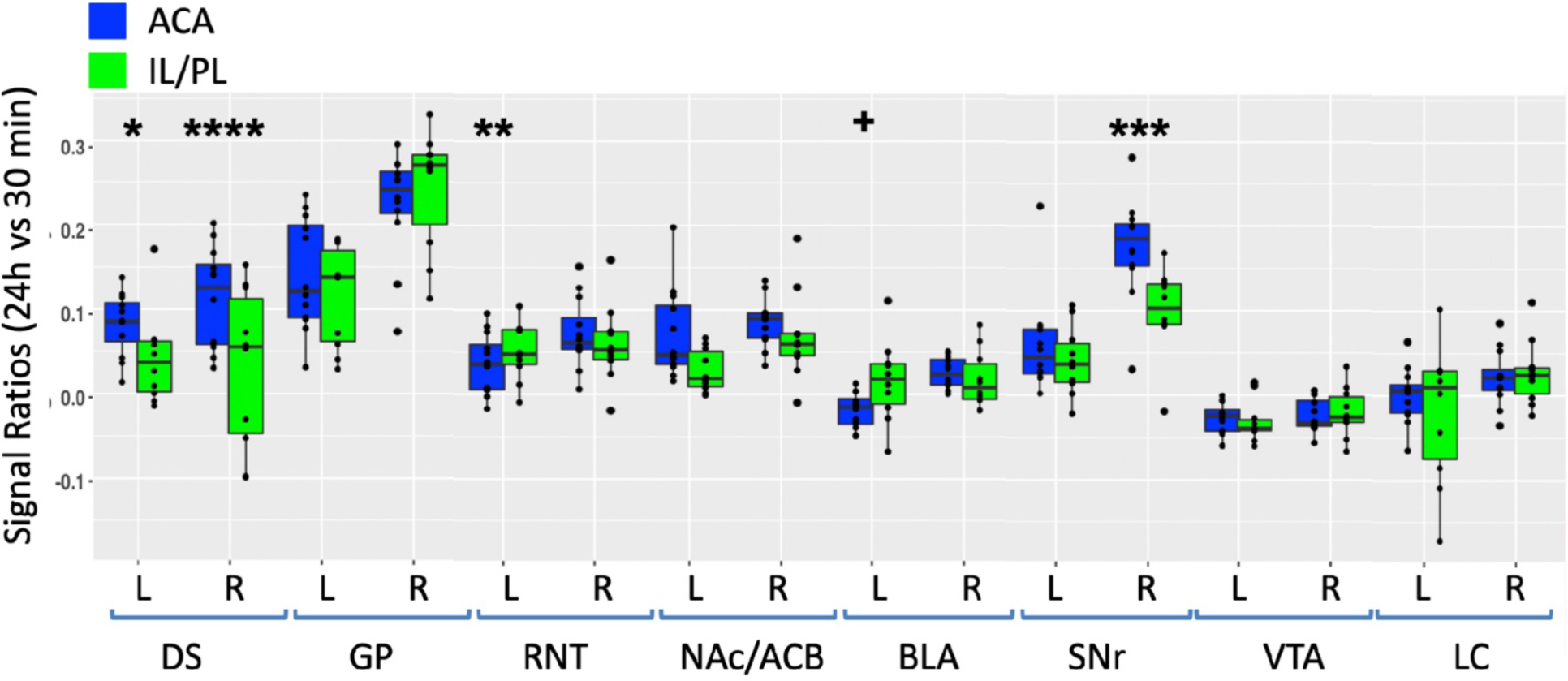
Between group comparisons of region of interest measurements of distal accumulations from either ACA and IL/PL injections. Box and whisker plots with individual values shown as scatter plots. Intensities were measured at 30m and 24h time points between the two groups in the same ROIs shown in Fig. 5, as indicated (Blue, ACA data; Green, IL/PL data). Ratios of 24h versus 30m were calculated and compared by linear mixed model. **^+^** p < 0.11; * p < 0.01; ** p < 0.001; *** p < 0.0001; **** p < 0.00001.

**Fig. 8.**
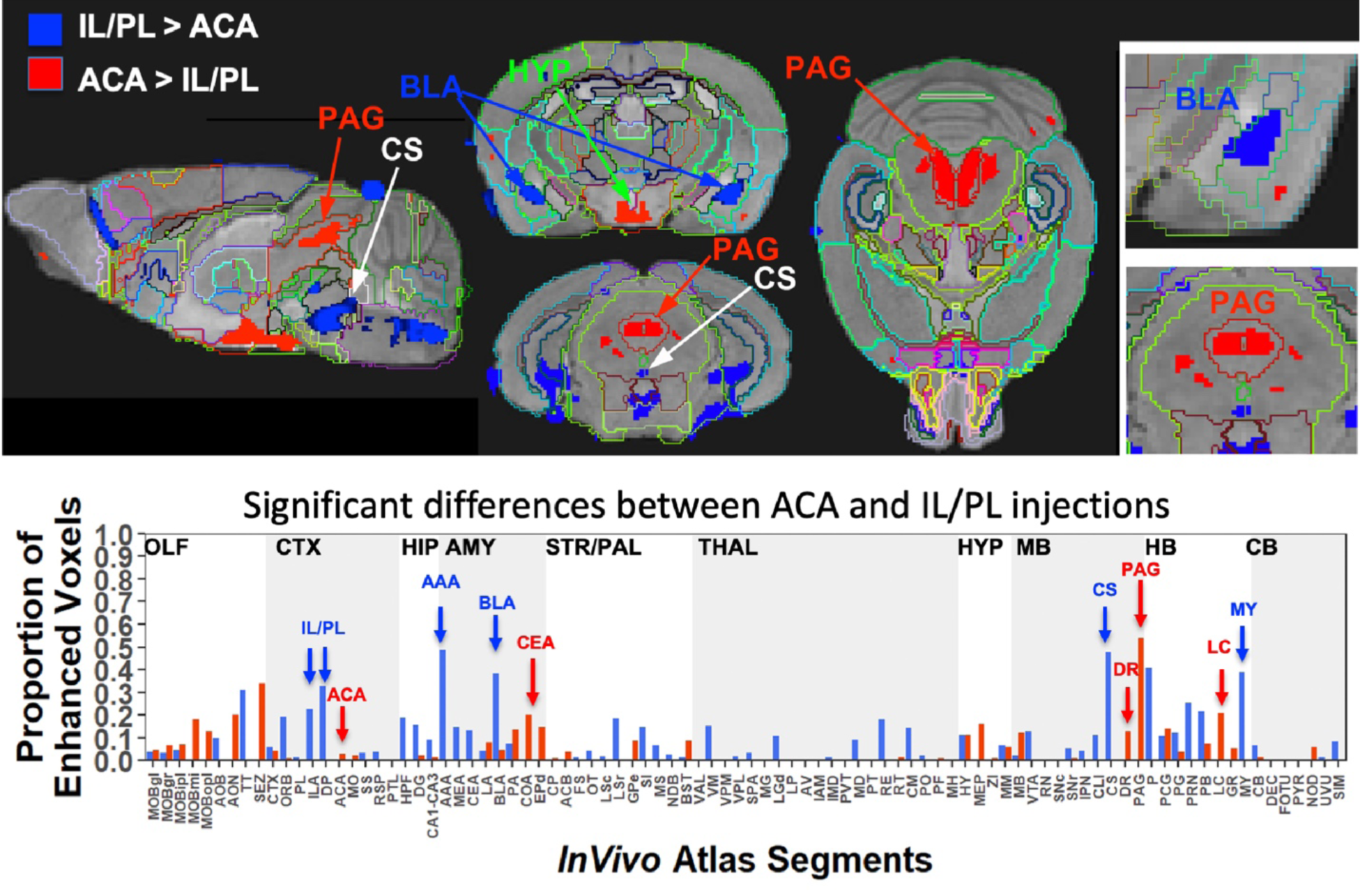
Between group statistical parametric mapping and segment-wise difference in volumes of Mn(II) accumulations. Two sample t-test between the two groups at 24h after injection in SPM produced two different maps (top panels). T-maps are overlaid on the grayscale MDT with *InVivo Atlas* superimposed as labels. Red, ACA greater than IL/PL, p < 0.001 FDR, T = 5.6 (red); and of IL/PL greater than ACA, p < 0.001 FDR, T = 5.1. Column graphs of fractional difference volumes (lower panel) identify major increases in the volumes of statistically significant voxels after ACA injections (Red) in the PAG, dorsal raphe (DR) and locus coeruleus (LC), and increases after IL/PL injections (Blue) in amygdala, particularly anterior amygdala area (AAA) and basolateral amygdala (BLA), as well as central superior raphe nucleus (CS) and medulla (MY) which is not extensively subdivided in this atlas. While statistically significant voxels increase in IL/PL in that group over the ACA group, only a minor increase in fraction of ACA enhanced voxels is found in the ACA group over the IL/PL. Because the high-concentration centroid of the injection site is less intense this could decrease the number of increased voxels found in the segment receiving the injection.

This SPM analysis was performed in two directions, with IL/PL greater than ACA injections and vice-versa. Resultant SPMs detect voxels with significant intensity differences between the two cohorts. Even at a fairly high stringency threshold of p < 0.001 FDR corrected (T = 5.6 for ACA and 5.1 for IL/PL), significantly different voxels within specific segments were highlighted. Segments with different volumes of significantly different intensity values were found within specific brain regions previously implicated in many of mPFC functions, such as amygdala, hypothalamus and monoaminergic systems. Increased volumes of Mn(II) accumulations occurred in different segments depending upon injection site (**Fig. 8, top panel**). By projecting both of these between group SPM maps at the same p value onto our *InVivo Atlas*, we found that segments with the largest volumes of significantly enhanced voxels that were also significantly different after IL/PL injections compared to those from ACA were basolateral amygdala (BLA) and central superior raphe (CS); whereas those less enhanced after IL/PL injection and more so after ACA injection were in the posterior hypothalamus (HYP) and periaqueductal gray (PAG) (**Fig. 8, lower panel** and **Table 2**). These images only represent significant differences between these two forebrain projection fields, they do not represent the full trajectory of projections from those two injection sites, as segments that are similar between them were not detected by this type of comparison. Analysis of the volumes of significant differences of statistically enhanced voxels per segment was performed and results visualized as column graphs (**Fig 8, lower panel**). In this case, volumes of enhanced voxels were extracted from the between-group unpaired two-tailed SPM t-maps, with threshold set at p < 0.01 FDR (ACA, T = 4.05; IL/PL, T = 3.6), lower stringency than the maps shown in the top panel. Many additional locations with 10-15% differences in active volumes were found. In small segments where even a small amount of signal would occupy a large ratio of voxels, this analysis matched visual inspection of the maps. However, in some other segments with larger number of total voxels, these calculations did not seem to reflect what was visible in the SPM maps. This issue was particularly troublesome in the hypothalamus, where localized differences between the two groups could be seen when scrolling through the 3D dataset but little difference appeared in the column graphs, and could affect other segments as well. Hence, we assumed that the large number of total voxels in the hypothalamic segment of our *InVivo Atlas* overwhelmed localized signal from differing sub-segments within the greater hypothalamus during ratiometric calculations.

**Table 2.** Differences in Projections from ACA and IL/PL. Anatomical differences are based on between-group two-tailed SPM t-test and correlations to our *InVivo Atlas*.

|  | ACA | IL/PL |
| --- | --- | --- |
| Domains | Segments | Segments |
| Amygdala | CEA | AAA |
|  | EPd | BLA |
| Midbrain | PAG | CS |
|  | DR |  |
| Hindbrain | LC | P (Pons NOS) |
|  |  | MY (Medulla NOS) |
| Hypothalamus | DVH | AHN |
|  | PVp | LHA |
|  | MOB |  |

To dive more deeply into possible differences between projections within the hypothalamus from these two sites, we sub-segmented this region on a high resolution, high contrast MEMRI image with anatomy based on grayscale contrast with reference to both the Allen Institute Mouse Brain Reference Atlas (Lein et al., 2007) and Paxinos (Vogt and Paxinos, 2014, Paxinos and Franklin, 2001) (**Fig. 9**), which are histologic atlases. Boundaries between smaller sub-domains in the hypothalamus can be difficult to define in the grayscale MR image even at high contrast and resolution. However, having an MR-based digital atlas aligned to our dataset greatly facilitated segment-wise analysis and was a major improvement over visual comparisons between MR images and histologic atlases. After inverse alignments of this sub-segmented atlas to our SPM maps, we could easily see dramatic differences in accumulations within specific identifiable sub-regions of the hypothalamus, any one of which could have major functional implications. The IL/PL injected cohort showed statistically significant accumulations in the anterior hypothalamic nucleus (AHN) and lateral hypothalamic area (LHA) (**Fig. 9, blue**) as well as BLA, previously recognized in **Fig. 8**, whereas the ACA (**Fig. 9, red**) had more accumulation than the IL/PL, with significantly different voxels clustering in the dorsomedial hypothalamus (DMH), periventricular hypothalamus posterior part (PVp) and mammillary body nuclei (MBO). While IL/PL projections accumulated Mn(II) signal more in anterior regions, at approximately bregma −0.7, ACA projections reached more caudally, approximately to bregma −1.48. Slice locations relative to bregma were determined by matching MR morphology with online Paxinos atlas (http://labs.gaidi.ca/mouse-brain-atlas/) (**Fig. 9, lower panels**). This represents a dramatic difference in average projection fields from these two closely placed forebrain regions, less than a millimeter apart anatomically, both in their targeting to PAG, a site of arousal, or to BLA, a site for fear processing, and to different sub-regions within the greater hypothalamus where hormonal responses are coupled to brain activity.

**Fig. 9.**
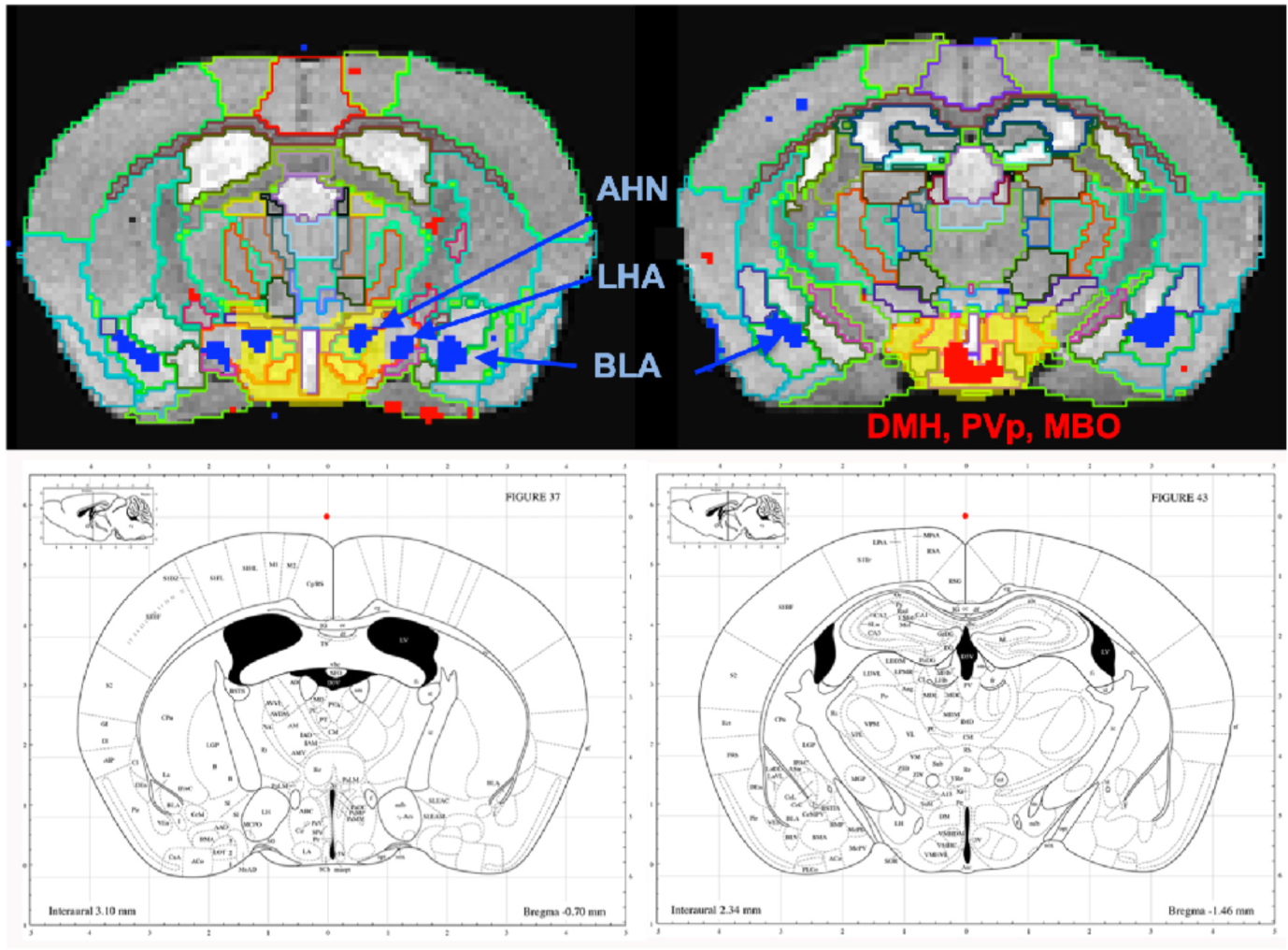
Differential accumulations of Mn(II) transported to the hypothalamus after IL/PL or ACA injections. Coronal slices at higher magnification of overlays of the two-sample t-test shown in F**ig. 8** onto a grayscale high contrast muse-Template grayscale image with sub-segmentation of the hypothalamus at coronal slices: A-P, 72 (IL/PL) and A-P, 81 (ACA), 900µm more posterior. The full hypothalamic segment is shown in yellow (top two panels). This segmentation was used to generate the segment-wise fractional difference volumes for the column graphs in **Fig. 8**. Here it is subdivided into anterior hypothalamic nuclei (AHN), lateral hypothalamic area (LHA), dorsal medial hypothalamus (DMH), periventricular hypothalamic nucleus, posterior part (PVp) and mammillary body (MBO), as indicated, with IL/PL (blue) and ACA (red). Screenshots of the online interactive Paxinos atlas at these slice positions (lower panel) (http://labs.gaidi.ca/mouse-brain-atlas/) at bregma −0.70 (left lower panel) and −1.49 (right lower panel).

## DISCUSSION

By employing MEMRI to trace projections from the medial prefrontal cortex (mPFC) in living animals after injection of tracer into two adjacent but distinct locations, we add to the growing literature about functional sub-domains within this region. Here we report that tracer injected into the ACA preferentially transports to posterior hypothalamic nuclei, mammillary nuclei, periaqueductal grey, central amygdalar nucleus, endopiriform nucleus, dorsal raphe and locus coeruleus as compared to the same tracer injected into the IL/PL, less than a millimeter away, which transports to more anterior segments in the hypothalamus, both anterior and basolateral amygdala, central raphe and various non-annotated discrete regions in pons and medulla. Thus, in mouse the two regions have distinct distal connections and may thus be capable of regulating varying limbic system responses.

This work is unique in a number of ways. By standardizing the injection sites, we traced projections from cohorts of 10-12 mice and report here average results across all animals in each cohort. No sacrifice was necessary, as transport was imaged in the living animal over a short, 24h, time period. By capturing images during transport and following progress in the living brain over time, we found that transport from a bolus of tracer could arrive at a distal destination within 6h, and may either increase into that target or dissipate. These data thus reveal that accurate distal projection mapping depends both on transport and clearance. Because we have an annotated digital atlas and a processing pipeline that allows us to average across all images in both cohorts with 22 mice and 3-4 images for each, our data is sufficient to gather statistically significant results. By measuring intensity values across all animals in candidate regions-of-interest, we obtain values for variance, average intensity changes, and statistically significance differences between time points both within the ACA group and between the two groups. This is in part made possible by the anatomical uniformity of the C57BL/6 mouse and by our computational intensity normalization and alignment procedures. The type of information derived from this study has never been available until now.

Some limitations of this study include differences between the two cohorts that may influence their forebrain projections, including gender mix and pre-injection history. Some evidence suggests that female gender affects mPFC (Knouse et al., 2022), although we have not found differences in previous work comparing MEMRI data between genders (Uselman et al., 2020). This result could have been due to a lower sensitivity to such gender-dependent effects in our methodology. Plasticity of medial forebrain projections has been hypothesized to play a role in drug use (Du et al., 2018, Frankin TR et al., 2002, Volkow et al., 1996) and in post-traumatic stress and anxiety disorders (Kesner and Churchwell, 2011, Laine et al., 2017, Padilla-Coreano et al., 2019, Sotres-Bayon and Quirk, 2010, Vertes, 2006). Thus, evidence exists that this circuit or network from mPFC into the limbic system may change in response to experience. It will be of interest to measure functional connections predicted here by more standard electrophysiological approaches. A next experiment would be to expose mice to cocaine and determine effects on these two projection fields in addicted versus naïve animals, possibly with MEMRI or other alternative mapping technology. It would be of interest if one or the other or both projection fields were redirected to alternate distal targets. Another experiment will be to test how early life adversity (ELA) affects forebrain projection anatomy in the adult, which would inform on developmental influences on mPFC control implicated in the known vulnerability after ELA to drug abuse and anxiety disorders.

Here Mn(II) distal accumulation serves as a proxy for axonal transport out axons to their distal, post-synaptic target, as we have done for hippocampal projections (Medina et al., 2019). We assume that this tracing is anterograde, occurs at similar rates between groups and is primarily functional, i.e. afferent projections that are electrically active. Mn(II) transport continues within axons in the absence of electrical activity. We previously found that Mn(II) does not accumulate in the superior colliculus in blind mice after injection in the eye despite entering intact retinal ganglion cells and traveling down their axons in the optic nerve (Bearer et al., 2007a). In sighted mice, Mn(II) crosses synapses in the superior colliculus, accumulating in post-synaptic neurons. Retrograde transport of Mn(II) has also been reported (Matsuda et al., 2010), in which case our data would detect both afferent and efferent connections between mPFC and deeper brain regions, although trans-synaptic tracing is unlikely in that scenario. While injection of MnCl_2_ at this concentration and volume appears safe in mouse, rat or even non-human primate (Uselman et al., 2022), there may be an acute excitatory event in neurons occasioned by a brief, locally high, concentration of cation. This effect could drive the Mn(II) across synapses not normally active at the time of tracing.

In conclusion, this study mapped brain-wide projections from ACA and IL/PL in living mice by MEMRI, and compared distal accumulations statistically. Significant differences were identified in key regions implicated in emotional states and their regulation, such as amygdala, hypothalamic subdomains and periaqueductal gray, as well as monoaminergic systems for serotonin and norepinephrine. Further experiments with electrophysiology will be needed to test these predicted downstream targets for their responses to medial forebrain activity. Next experiments with MEMRI should focus on the plasticity of these projections in response to experience and/or drug use.

## ACKNOWLEDGEMENTS

We are grateful to the Beckman Institute of California Institute of Technology for use of the 11.7 Bruker scanner and support of the Caltech Biological Imaging Facility. For technical support, we thank Sharon Wu Lin for animal handling, Xiaowei Zhang for scanner operation and image capture, Adam W. Mitchell, a UNM computer science graduate student for the initial steps in processing the IL/PL images, James Chavez for IT support, and Kathleen Kilpatrick, Kevin Reagan, Daniel R. Perez for laboratory management. Funded by NIDA R01 DA018184 (REJ), NIMH R01MH096093 (ELB) and the Harvey Family Endowment to UNM Foundation (ELB).

## Notes

### Competing Interest Statement

The authors have declared no competing interest.

## References

1. Ashburner, J., Barnes, G., Chen, C., Daunizeau, J., Flandin, G., Friston, K., Gitelman, D., Kiebel, S., Kilner, J. & Litvak, V. J. F. I. L., INSTITUTE OF NEUROLOGY 2012. SPM8 manual.

2. Ashburner, J., Barnes, G., Chen, C.-C., Daunizeau, J., Flandin, G., Friston, K., Kiebel, S., Kilner, J., Litvak, V. & Moran, R. J. W. T. C. F. N., LONDON, UK 2014. SPM12 manual. 2464.

3. Aydogan, D. B., Jacobs, R., Dulawa, S., Thompson, S. L., Francois, M. C., Toga, A. W., Dong, H., Knowles, J. A. & Shi, Y. 2018. When tractography meets tracer injections: a systematic study of trends and variation sources of diffusion-based connectivity. Brain Struct Funct, 223, 2841–2858.

4. Barron, H. C., Mars, R. B., Dupret, D., Lerch, J. P. & Sampaio-Baptista, C. 2021. Cross-species neuroscience: closing the explanatory gap. Philosophical Transactions of the Royal Society B, 376, 20190633.

5. Bearer, E. L., Falzone, T. L., Zhang, X., Biris, O., Rasin, A. & Jacobs, R. E. 2007a. Role of neuronal activity and kinesin on tract tracing by manganese-enhanced MRI (MEMRI). Neuroimage, 37 Suppl 1, S37–46.

6. Bearer, E. L., Manifold-Wheeler, B. C., Medina, C. S., Gonzales, A. G., Chaves, F. L. & Jacobs, R. E. 2018. Alterations of functional circuitry in aging brain and the impact of mutated APP expression. Neurobiol Aging, 70, 276–290.

7. Bearer, E. L. & Wu, C. 2019. Herpes Simplex Virus, Alzheimer’s Disease and a Possible Role for Rab GTPases. Frontiers in Cell and Developmental Biology, 7.

8. Bearer, E. L., Zhang, X., Biris, O. & Jacobs, R. E. 2009a. Imaging biophysics of axonal transport with MEMRI: Optic tract transport is altered in mouse model of Alzheimer’s disease. Proc Int Soc Magn Reson Med Sci Meet Exhib Int Soc Magn Reson Med Sci Meet Exhib, 2009, 540.

9. Bearer, E. L., Zhang, X. & Jacobs, R. E. 2007b. Live imaging of neuronal connections by magnetic resonance: Robust transport in the hippocampal-septal memory circuit in a mouse model of Down syndrome. Neuroimage, 37, 230–42.

10. Bearer, E. L., Zhang, X. & Jacobs, R. E. 2022. Studying Axonal Transport in the Brain by Manganese-Enhanced Magnetic Resonance Imaging). In: Vagnoni, A. (ed.) Axonal Transport: Methods and Protocols. New York, NY: Springer US.

11. Bearer, E. L., Zhang, X., Janvelyan, D., Boulat, B. & Jacobs, R. E. 2009b. Reward circuitry is perturbed in the absence of the serotonin transporter. Neuroimage, 46, 1091–104.

12. Bedenk, B. T., Almeida-Correa, S., Jurik, A., Dedic, N., Grunecker, B., Genewsky, A. J., Kaltwasser, S. F., Riebe, C. J., Deussing, J. M., Czisch, M. & Wotjak, C. T. 2018. Mn(2+) dynamics in manganese-enhanced Mri (MEMRI): Cav1.2 channel-mediated uptake and preferential accumulation in projection terminals. Neuroimage, 169, 374–382.

13. Bubb, E. J., Metzler-Baddeley, C. & Aggleton, J. P. 2018. The cingulum bundle: Anatomy, function, and dysfunction. Neurosci Biobehav Rev, 92, 104–127.

14. Cox, R. W. 1996. Afni: Software for Analysis and Visualization of Functional Magnetic Resonance Neuroimages. Computers and Biomedical Research, 29, 162–173.

15. Damasio, H., Grabowski, T., Frank, R., Galaburda, A. M. & Damasio, A. R. 1994. The Return of Phineas Gage: Clues About the Brain from the Skull of a Famous Patient. Science, 264, 1102–1105.

16. Delora, A., Gonzales, A., Medina, C. S., Mitchell, A., Mohed, A. F., Jacobs, R. E. & Bearer, E. L. 2016. A simple rapid process for semi-automated brain extraction from magnetic resonance images of the whole mouse head. J Neurosci Methods, 257, 185–93.

17. Du, C., Volkow, N. D., You, J., Park, K., Allen, C. P., Koob, G. F. & Pan, Y. 2018. Cocaine-induced ischemia in prefrontal cortex is associated with escalation of cocaine intake in rodents. Molecular Psychiatry, 25, 1759–1776.

18. Elluru, R. G., Bloom, G. S. & Brady, S. T. 1995. Fast axonal transport of kinesin in the rat visual system: functionality of kinesin heavy chain isoforms. Molecular Biology of the Cell, 6, 21–40.

19. Fillinger, C., Yalcin, I., Barrot, M. & Veinante, P. 2018. Efferents of anterior cingulate areas 24a and 24b and midcingulate areas 24a’ and 24b’ in the mouse. Brain Struct Funct, 223, 1747–1778.

20. Frankin Tr, Acton Pd, Maldjian Ja, Gray Jd, Crofit Jr, Dackis Ca, O’brien Cp & Ar, C. 2002. Decreased Gray Matter Concentration in the Insular,Orbitofrontal, Cingulate, and Temporal Cortices ofCocaine Patients. Biol Psychiatry, 51, 134–162.

21. Friston, K. J. 1996. Statistical parametric mapping and other analysis of functional imaging data. In: Toga, A. W. & Mazziotta, J. C. (eds.) Brain Mapping: The Methods. San Diego: Academic Press.

22. Gabbott, P. L. A., Warner, T. A., Jays, P. R. L., Salway, P. & Busby, S. J. 2005. Prefrontal cortex in the rat: Projections to subcortical autonomic, motor, and limbic centers. The Journal of Comparative Neurology, 492, 145–177.

23. Gallagher, J. J., Zhang, X., Hall, F. S., Uhl, G. R., Bearer, E. L. & Jacobs, R. E. 2013. Altered reward circuitry in the norepinephrine transporter knockout mouse. PLos One, 8, e57597.

24. Gallagher, J. J., Zhang, X., Ziomek, G. J., Jacobs, R. E. & Bearer, E. L. 2012. Deficits in axonal transport in hippocampal-based circuitry and the visual pathway in APP knock-out animals witnessed by manganese enhanced MRI. Neuroimage, 60, 1856–66.

25. Jenkinson, M., Bannister, P., Brady, M. & Smith, S. 2002. Improved optimization for the robust and accurate linear registration and motion correction of brain images. Neuroimage, 17, 825–41.

26. Jenkinson, M., Beckmann, C. F., Behrens, T. E., Woolrich, M. W. & Smith, S. M. 2012a. FSL. Neuroimage, 62, 782–90.

27. Jenkinson, M., Beckmann, C. F., Behrens, T. E., Woolrich, M. W. & Smith, S. M. J. N. 2012b. Fsl. 62, 782–790.

28. Jing, D., Zhang, S., Luo, W., Gao, X., Men, Y., Ma, C., Liu, X., Yi, Y., Bugde, A., Zhou, B. O., Zhao, Z., Yuan, Q., Feng, J. Q., Gao, L., Ge, W. P. & Zhao, H. 2018. Tissue clearing of both hard and soft tissue organs with the PEGASOS method. Cell Res, 28, 803–818.

29. Kesner, R. P. & Churchwell, J. C. 2011. An analysis of rat prefrontal cortex in mediating executive function. Neurobiology of Learning and Memory, 96, 417–431.

30. Knouse, M. C., Mcgrath, A. G., Deutschmann, A. U., Rich, M. T., Zallar, L. J., Rajadhyaksha, A. M. & Briand, L. A. 2022. Sex differences in the medial prefrontal cortical glutamate system. Biol Sex Differ, 13, 66.

31. Laine, M. A., Sokolowska, E., Dudek, M., Callan, S. A., Hyytia, P. & Hovatta, I. 2017. Brain activation induced by chronic psychosocial stress in mice. Sci Rep, 7, 15061.

32. Lein, E. S., Hawrylycz, M. J., Ao, N., Ayres, M., Bensinger, A., Bernard, A., Boe, A. F., Boguski, M. S., Brockway, K. S., Byrnes, E. J., Chen, L., Chen, L., Chen, T. M., Chin, M. C., Chong, J., Crook, B. E., Czaplinska, A., Dang, C. N., Datta, S., Dee, N. R., Desaki, A. L., Desta, T., Diep, E., Dolbeare, T. A., Donelan, M. J., Dong, H. W., Dougherty, J. G., Duncan, B. J., Ebbert, A. J., Eichele, G., Estin, L. K., Faber, C., Facer, B. A., Fields, R., Fischer, S. R., Fliss, T. P., Frensley, C., Gates, S. N., Glattfelder, K. J., Halverson, K. R., Hart, M. R., Hohmann, J. G., Howell, M. P., Jeung, D. P., Johnson, R. A., Karr, P. T., Kawal, R., Kidney, J. M., Knapik, R. H., Kuan, C. L., Lake, J. H., Laramee, A. R., Larsen, K. D., Lau, C., Lemon, T. A., Liang, A. J., Liu, Y., Luong, L. T., Michaels, J., Morgan, J. J., Morgan, R. J., Mortrud, M. T., Mosqueda, N. F., Ng, L. L., Ng, R., Orta, G. J., Overly, C. C., Pak, T. H., Parry, S. E., Pathak, S. D., Pearson, O. C., Puchalski, R. B., Riley, Z. L., Rockett, H. R., Rowland, S. A., Royall, J. J., Ruiz, M. J., Sarno, N. R., Schaffnit, K., Shapovalova, N. V., Sivisay, T., Slaughterbeck, C. R., Smith, S. C., Smith, K. A., Smith, B. I., Sodt, A. J., Stewart, N. N., Stumpf, K. R., Sunkin, S. M., Sutram, M., Tam, A., Teemer, C. D., Thaller, C., Thompson, C. L., Varnam, L. R., Visel, A., Whitlock, R. M., Wohnoutka, P. E., Wolkey, C. K., Wong, V. Y., et al. 2007. Genome-wide atlas of gene expression in the adult mouse brain. Nature, 445, 168–76.

33. Matsuda, K., Wang, H. X., Suo, C., Mccombe, D., Horne, M. K., Morrison, W. A. & Egan, G. F. 2010. Retrograde axonal tracing using manganese enhanced magnetic resonance imaging. Neuroimage, 50, 366–74.

34. Mcauliffe, M. J., Lalonde, F. M., Mcgarry, D., Gandler, W., Csaky, K. & Trus, B. L. Medical Image Processing, Analysis and Visualization in clinical research. Proceedings 14th IEEE Symposium on Computer-Based Medical Systems. CBMS 2001, 26-27 July 2001 2001. 381–386.

35. Medina, C. S., Biris, O., Falzone, T. L., Zhang, X., Zimmerman, A. J. & Bearer, E. L. 2017a. Hippocampal to basal forebrain transport of Mn2+ is impaired by deletion of KLC1, a subunit of the conventional kinesin microtubule-based motor. Neuroimage, 145, 44–57.

36. Medina, C. S., Manifold-Wheeler, B., Gonzales, A. & Bearer, E. L. 2017b. Automated Computational Processing of 3-D MR Images of Mouse Brain for Phenotyping of Living Animals. Curr Protoc Mol Biol, 119, 29a 5 1–29a 5 38.

37. Medina, C. S., Uselman, T. W., Barto, D. R., Chaves, F., Jacobs, R. E. & Bearer, E. L. 2019. Decoupling the Effects of the Amyloid Precursor Protein From Amyloid-beta Plaques on Axonal Transport Dynamics in the Living Brain. Front Cell Neurosci, 13, 501.

38. Narita, K., Kawasaki, F. & Kita, H. 1990. Mn and Mg influxes through Ca channels of motor nerve terminals are prevented by verapamil in frogs. Brain Res, 510, 289–95.

39. Padilla-Coreano, N., Canetta, S., Mikofsky, R. M., Alway, E., Passecker, J., Myroshnychenko, M. V., Garcia-Garcia, A. L., Warren, R., Teboul, E., Blackman, D. R., Morton, M. P., Hupalo, S., Tye, K. M., Kellendonk, C., Kupferschmidt, D. A. & Gordon, J. A. 2019. Hippocampal-Prefrontal Theta Transmission Regulates Avoidance Behavior. Neuron, 104, 601–610.e4.

40. Pautler, R. G. 2006. Biological applications of manganese-enhanced magnetic resonance imaging. Methods Mol Med, 124, 365–86.

41. Pautler, R. G. & Koretsky, A. P. 2002. Tracing odor-induced activation in the olfactory bulbs of mice using manganese-enhanced magnetic resonance imaging. Neuroimage, 16, 441–8.

42. Pautler, R. G., Mongeau, R. & Jacobs, R. E. 2003. In vivo trans-synaptic tract tracing from the murine striatum and amygdala utilizing manganese enhanced Mri (MEMRI). Magnetic Resonance in Medicine, 50, 33–39.

43. Paxinos, G. & Franklin, K. 2001. The Mouse Brain in Stereotaxic Coordinates, San Diego, Academic Press.

44. Riga, D., Matos, M. R., Glas, A., Smit, A. B., Spijker, S. & Van Den Oever, M. C. 2014. Optogenetic dissection of medial prefrontal cortex circuitry. Frontiers in Systems Neuroscience, 8.

45. Satpute-Krishnan, P., Degiorgis, J. A., Conley, M. P., Jang, M. & Bearer, E. L. 2006. A peptide zipcode sufficient for anterograde transport within amyloid precursor protein. Proceedings of the National Academy of Sciences, 103, 16532–16537.

46. Sesack, S. R. & Pickel, V. M. 1992. Prefrontal cortical efferents in the rat synapse on unlabeled neuronal targets of catecholamine terminals in the nucleus accumbens septi and on dopamine neurons in the ventral tegmental area. J Comp Neurol, 320, 145–60.

47. Sled, J. G., Zijdenbos, A. P. & Evans, A. C. 1998. A nonparametric method for automatic correction of intensity nonuniformity in MRI data. IEEE Trans Med Imaging, 17, 87–97.

48. Smith, S. M., Jenkinson, M., Woolrich, M. W., Beckmann, C. F., Behrens, T. E., Johansen-Berg, H., Bannister, P. R., De Luca, M., Drobnjak, I., Flitney, D. E., Niazy, R. K., Saunders, J., Vickers, J., Zhang, Y., De Stefano, N., Brady, J. M. & Matthews, P. M. 2004. Advances in functional and structural MR image analysis and implementation as FSL. Neuroimage, 23 Suppl 1, S208–19.

49. Sotres-Bayon, F. & Quirk, G. J. 2010. Prefrontal control of fear: more than just extinction. Curr Opin Neurobiol, 20, 231–5.

50. Tustison, N. J., Avants, B. B., Cook, P. A., Zheng, Y., Egan, A., Yushkevich, P. A. & Gee, J. C. 2010. N4itk: Improved N3 Bias Correction. IEEE Transactions on Medical Imaging, 29, 1310–1320.

51. Uselman, T. W., Barto, D. R., Jacobs, R. E. & Bearer, E. L. 2020. Evolution of brain-wide activity in the awake behaving mouse after acute fear by longitudinal manganese-enhanced MRI. Neuroimage, 222, 1–22.

52. Uselman, T. W., Medina, C. S., Gray, H. B., Jacobs, R. E. & Bearer, E. L. 2022. Longitudinal manganese-enhanced magnetic resonance imaging of neural projections and activity. NMR Biomed, 35, e4675.

53. Van Heukelum, S., Mars, R. B., Guthrie, M., Buitelaar, J. K., Beckmann, C. F., Tiesinga, P. H. E., Vogt, B. A., Glennon, J. C. & Havenith, M. N. 2020. Where is Cingulate Cortex? A Cross-Species View. Trends Neurosci, 43, 285–299.

54. Vertes, R. P. 2006. Interactions among the medial prefrontal cortex, hippocampus and midline thalamus in emotional and cognitive processing in the rat. Neuroscience, 142, 1–20.

55. Vogt, B. A. & Paxinos, G. 2014. Cytoarchitecture of mouse and rat cingulate cortex with human homologies. Brain Struct Funct, 219, 185–92.

56. Volkow, N. D., Ding, Y. S., Fowler, J. S. & Wang, G. J. 1996. Cocaine addiction: hypothesis derived from imaging studies with Pet. J Addict Dis, 15, 55–71.

57. Watanabe, T., Natt, O., Boretius, S., Frahm, J. & Michaelis, T. 2002. In vivo 3d MRI staining of mouse brain after subcutaneous application of MnCl2. Magnetic Resonance in Medicine, 48, 852–859.

58. Zhang, X., Bearer, E. L., Boulat, B., Hall, F. S., Uhl, G. R. & Jacobs, R. E. 2010. Altered neurocircuitry in the dopamine transporter knockout mouse brain. PLos One, 5, e11506.

59. Zingg, B., Hintiryan, H., Gou, L., Song, M. Y., Bay, M., Bienkowski, M. S., Foster, N. N., Yamashita, S., Bowman, I., Toga, A. W. & Dong, H. W. 2014. Neural networks of the mouse neocortex. Cell, 156, 1096–111.

