## Supplemental Materials for "Harnessing axonal transport to map reward circuitry: Differing brain-wide projections from medial forebrain domains"

### 1 Supplementary Data List

Table S1. Abbreviations (pdf)
Table S2. ROI coordinates (pdf
Table S3. ROI Measurements (excel)
Table S4. ROI Statistics (pdf)

Supplemental Fig. S1. Behavior
Supplemental Fig. S2. Injection site overlays

**Supplementary Table S1**. Abbreviations used in the column graphs in Fig. 6 and 8, in the order in which the columns appear in the graph. Nomenclature is according to the Allen Institute for Brain Science Mouse Brain Reference Atlas.

| Abbreviation | Nominal Label | Abbreviation | Nominal Label |
|---|---|---|---|
| AAA | Anterior amygdalar area | MO | Somatomotor areas |
| ACA | Anterior cingulate area | MOBgl | Main olfactory bulb glomerular |
| ACB | Nucleus accumbens (a.k., NAc) | MOBgr | Main olfactory bulb granule layer |
| aco | Anterior commissure olfactory limb | MOBipl | Main olfactory bulb inner plexiform layer |
| AOB | Accessory olfactory bulb | MOBmi | Main olfactory bulb mitral layer |
| AON | Anterior olfactory nucleus | MOBopl | Main olfactory bulb outer plexiform layer |
| aot | Accessory optic tract | MS | Medial septal nucleus |
| AV | Anteroventral nucleus of thalamus | MY | Medulla |
| BLA | Basolateral amygdala nucleus | NDB | Diagonal band nucleus |
| BST | Bed nuclei of the stria terminalis | NOD | Nodulus X |
| CA1-CA3 | Field ca1 ca2 ca3 pyramidal layer | onl | Nerve layer of main olfactory bulb |
| CB | Cerebellum | opt | Optic tract |
| cc | Corpus callosum | ORB | Orbital area |
| CEA | Central amygdalar nucleus | OT | Olfactory tubercle |
| CLI | Central linear nucleus raphe | P | Pons |
| CM | Central medial nucleus of the thalamus | PA | Posterior amygdalar nucleus |
| COA | Cortical amygdala area | PAG | Periaqueductal gray |
| CP | Caudoputamen | PB | Parabrachial nucleus |
| CS | Superior central raphe nucleus | PCG | Pontine central gray |
| CTX | Cerebral cortex | PF | Parafascicular nucleus |
| DEC | Declive VI | PG | Pontine gray |
| DG | Dentate gyrus | PL | Prelimbic area |
| DP | Dorsal peduncular area | PO | Posterior complex of the thalamus |
| DR | Dorsal raphe nucleus | PRN | Pontine reticular nucleus |
| em | External medullary lamina | PT | Parataenial nucleus |
| EPd | Endopiriform nucleus dorsal part | PTL | Posterior parietal association areas |
| fi | Fimbria | PVT | Paraventricular nucleus of the thalamus |
| FOTU | Folium-tuber Vermis VII | PYR | Pyramus VIII |
| FS | Fundus of striatum | RE | Nucleus of reunions |
| GPe | Globus pallidus | RN | Red nucleus |
| GR | Gracile nucleus | RSP | Retrosplenial area |
| HPF | Hippocampal formation | RT | Reticular nucleus of the thalamus |
| HY | Hypothalamus | SEZ | Subependymal zone |
| IAM | Interanteromedial nucleus of thalamus | SI | Substantia innominata |
| ILA | Infralimbic area | SIM | Simple lobule |
| IMD | Intermedial dorsal thalamus | sm | Stria medullaris |
| int | Internal capsule | SNc | Substantia nigra compact part |
| IPN | Interpeduncular nucleus | SNr | Substantia nigra reticular part |
| LA | Lateral amygdala nucleus | SPA | Subparafascicular area |
| LC | Loc coeruleus | SS | Somatosensory areas |
| LGd | Lateral geniculate complex dorsal part | st | Stria terminalis |
| lot | Lateral olfactory tract body | TT | Taenia tecta dorsal part |
| LP | Lateral posterior nucleus of the thalamus | UVU | Uvula IX |
| LSc | Lateral septal nucleus caudal part | V3 | Third ventricle |
| LSr | Lateral septal nucleus rostral part | VAL | Ventral anterior lateral thalamic complex |
| MB | Midbrain | vhc | Ventral hippocampal commissure |
| MD | Mediodorsal nucleus of thalamus | VL | Lateral ventricle |
| MEA | Medial amygdalar area | VM | Ventral posterolateral thalamus |
| MEP | Median preoptic nucleus | VPL | Ventral posteromedial nucleus of the thalamus |
| MG | Medial geniculate complex | VPM | Ventral medial thalamic nucleus |
| MH | Medial habenula | VTA | Ventral tegmental area |
| MM | Medial mammillary nucleus | ZI | Zona incerta |

**Supplementary Table S2.** Coordinates for Regions of Interest Analysis relative to Bregma (in mm)

| **Region of Interest** | **Hemisphere** | **Coordinates** | | |
|---|---|---|---|---|
| | | **ML** | **DV** | **AP** |
| DS | Left | -1.5 | 0.8 | 2.7 |
| | Right | 1.4 | 0.8 | 3.7 |
| GP | Left | -1.6 | -0.3 | 4.7 |
| | Right | 1.7 | -0.3 | 4.7 |
| RNT | Left | -1.2 | -0.5 | 4.2 |
| | Right | 1.3 | -1.0 | 4.1 |
| NAc/ACB | Left | -1.0 | 1.0 | 5.5 |
| | Right | 1.2 | 1.0 | 5.4 |
| BLA | Left | -3.1 | -2.0 | 5.9 |
| | Right | 3.6 | -2.0 | 5.6 |
| SNr | Left | -1.3 | -3.1 | 5.4 |
| | Right | 1.5 | -3.0 | 5.4 |
| VTA | Left | -0.8 | -3.0 | 5.2 |
| | Right | 0.7 | -3.0 | 5.1 |
| LC | Left | -0.9 | -5.7 | 4.4 |
| | Right | 0.8 | -5.7 | 4.6 |

Bregma locations corresponding to FSL (*fslroi*) defined 3 x 3 x 3 voxel cubes.

**Supplementary Table S3. ROI Measurements**

**See Supplementary Table S3, a separate file available as Excel.**

**Supplementary Table S4. ROI Statistics (pdf)**

Supplementary Table S4. Statistics for ROI within and between group analyses Figures 5 and 7 in the Main text.

**Statistics for ACA within group analysis for differences at 6h vs 24h**

| **ROI** | **t.ratio** | **p value** | **Asterisks** |
|---|---|---|---|
| DS-L2 | 3.289 | 0.0006 | *** <0.001 |
| GP_L | -2.861 | 0.0045 | ** <0.01 |
| GP_R | -4.304 | <0.0001 | **** |
| NAC_R2 | -1.682 | 0.0936 | * <0.1 |
| RNT_L2 | 3.212 | 0.0014 | * <0.01 |
| RNT_R2 | 4.468 | <0.0001 | **** |
| SNR_L | -2.788 | 0.0056 | ** <0.01 |
| SNR_R | -5.578 | <0.0001 | **** |
| LC_L | 3.124 | 0.0019 | ** <0.01 |
| LC_R | 2.680 | 0.0077 | ** <0.01 |

**Statistics for between ACA and IL/PL cohorts at 24h post-injection**

| Region | t.ratio | p value | Asterisks |
|---|---|---|---|
| DS_L2 | 0.39 | 0.0642 | * <0.1 |
| DS_R2 | 3.917 | 0.0009 | **** <0.001 |
| RNT_L2 | 2.1 | 0.0486 | ** <0.05 |
| BLA_L | -1.692 | 0.106 | + <= 0.2 |
| SNr_R | 3.624 | 0.0017 | ***<0.005 |

### 2.2 Supplementary Figures

**Supplementary Figure S1: Time spent not moving**

Mice were video recorded during the last 10m of 30m time spent in a custom arena at two timepoints before the forebrain injections: At baseline before any handling and at 23 days after handling, imaging and housing. Time spent not-moving within each 1-minute interval was tabulated in Ethovision and results graphed in Excel (Microsoft Office). Statistical comparisons were performed in R by ANOVA between these two time points. A small but statistically significant difference was found between baseline and 23d ($p < 0.01$).

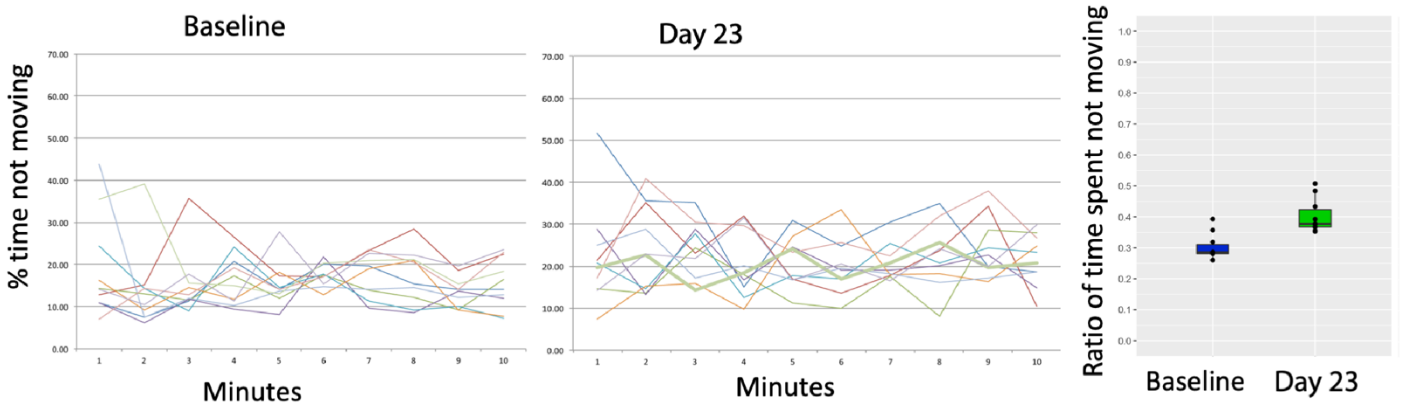

**Supplementary Figure S2**. Statistical maps of ACA and IL/PL injection sites

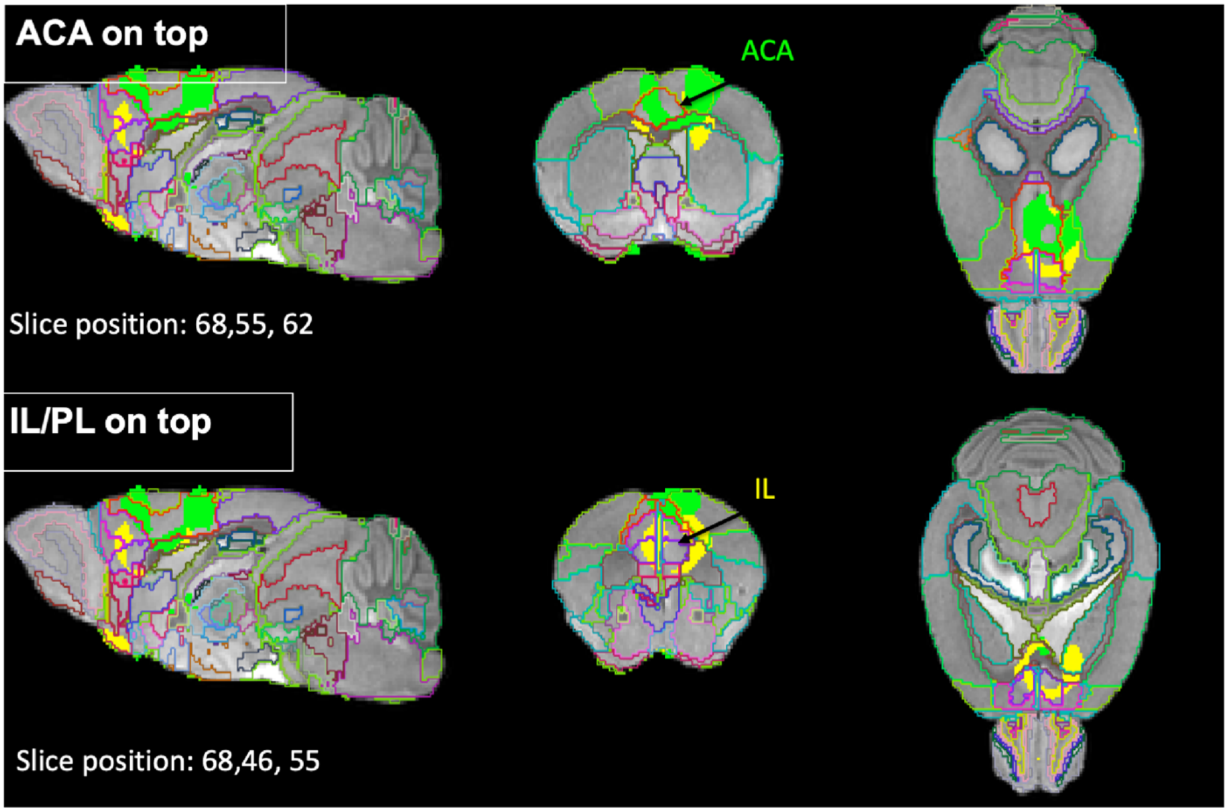


The 30m post-injection images for each cohort were compared to pre-injection image by a within-group paired t-test in SPM. Resultant maps of significantly enhanced voxels at a threshold of $p < 0.05$ FDR corrected (T values: IL/PL, T = 4.75; ACA, T = 4.05) were overlaid on the template image and with *InVivo Atlas* v.10 on the last layer to define position. Slice positions are indicated as voxel positions in the 3D dataset. Note that slice position of the injection halo representing Mn(II) diffusion out of the injection site differs in AP dimension by 9 voxels (0.9mm) and in the DV dimension by 7 voxels (0.7mm), and that there is some overlap in the region between the two sites. Bright pink outline delineates the ACA, and purple the Infralimbic segments as indicated on the coronal slices.
